# The Anti-*Burkholderia* Lasso Peptide Ubonodin Co-Opts the Siderophore Receptor PupB for Cellular Entry

**DOI:** 10.1101/2022.03.09.483586

**Authors:** Truc Do, Alina Thokkadam, Robert Leach, A. James Link

**Affiliations:** Department of Chemical and Biological Engineering, Princeton University, Princeton, NJ 08544, United States; Lewis-Sigler Institute for Integrative Genomics, Princeton University, Princeton, NJ 08544, United States; Department of Chemistry, Princeton University, Princeton, NJ 08544, United States; Department of Molecular Biology, Princeton University, Princeton, NJ 08544, United States

## Abstract

New antibiotics are needed as bacterial infections continue to be a leading cause of death. Notorious among antibiotic-resistant bacteria is the Burkholderia *cepacia* complex (Bcc), which infects cystic fibrosis patients, causing lung function decline. We recently discovered a novel ribosomally synthesized and post-translationally modified peptide (RiPP), ubonodin, with potent activity against several *Burkholderia* pathogens. Ubonodin inhibits RNA polymerase, but only select Bcc strains were susceptible, indicating that having a conserved cellular target does not guarantee activity. Given the cytoplasmic target, we speculate that cellular uptake of ubonodin determines susceptibility. Here, we report a new outer membrane siderophore receptor, PupB, that is required for ubonodin uptake in *B. cepacia*. Loss of PupB renders *B. cepacia* resistant to ubonodin, whereas expressing PupB sensitizes a resistant strain. Thus, outer membrane transport is the major determinant of ubonodin’s spectrum of activity. We also show that PupB is activated by a TonB protein and examine a transcriptional pathway that further regulates PupB. Finally, we elucidate the complete cellular uptake pathway for ubonodin by also identifying its inner membrane transporter in *B. cepacia*. Our work unravels central steps in the mechanism of action of ubonodin and establishes a general framework for dissecting RiPP function.

## INTRODUCTION

The *Burkholderia cepacia* complex (Bcc) is a group of Gram-negative bacteria that can cause serious, often fatal infections in individuals living with cystic fibrosis or other underlying pulmonary disease (Chiarini et al., 2006; Leitão et al., 2017; Mahenthiralingam et al., 2005). Unfortunately, only a few antibiotics are effective against opportunistic Bcc pathogens due to their high level of intrinsic resistance, so there is a need to develop new anti-Bcc compounds and decipher their mechanisms of action. Throughout history, humans have looked to nature as a rich and abundant source of antimicrobial compounds. Notable among antimicrobial natural products are ribosomally synthesized and post-translationally modified peptides (RiPPs). RiPPs are synthesized as gene-encoded precursor peptides that are post-translationally tailored with additional chemical modifications to form the mature peptide scaffold (Arnison et al., 2012; Montalbán-López et al., 2020). Lasso peptides are a growing class of bacterial-derived RiPPs that are characterized by their unique lariat knot conformation, formed by the C-terminal tail threading through a macrolactam ring (**Figure 1A**) (Hegemann et al., 2015; Maksimov et al., 2012). The ring is forged by connecting the N-terminus of the peptide to an acidic side chain via an isopeptide bond. The loop and tail are held in place by bulky steric lock residues that straddle the ring or by disulfide bridges. Despite a relatively conserved three-dimensional structure, lasso peptides are remarkably diverse in biological function (Cao et al., 2021; Kodani and Unno, 2020; Li and Rebuffat, 2020). However, as with most RiPPs, the mechanisms of action of lasso peptides are poorly characterized, hampering efforts to develop them as drugs.

**Figure 1.**
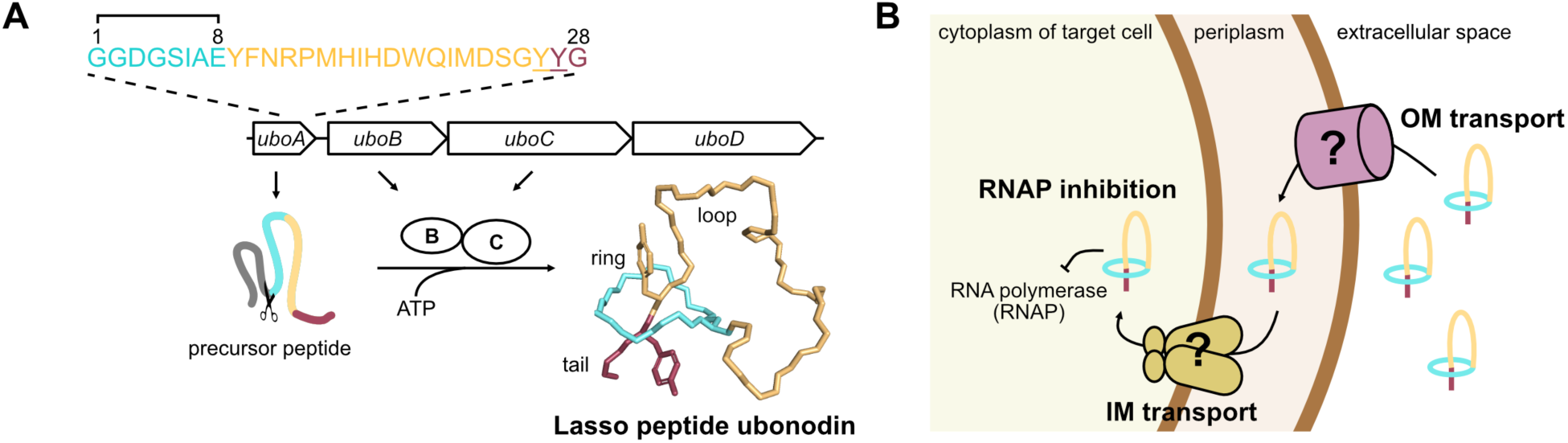
Biosynthesis and mechanism of action of ubonodin. (**A**) Schematic showing the ubonodin biosynthetic gene cluster (BGC). Following production of the ubonodin precursor peptide specified by *uboA*, the B enzyme cleaves the leader peptide (gray), generating the core peptide whose amino acid sequence is outlined at the top. The C enzyme forms the final lariat structure by connecting the amino group of Gly1 to the Glu8 side chain, forging an 8-membered ring through which the tail is threaded. Residues Tyr26 and Tyr27 (underlined in the sequence) act as steric locks to prevent ubonodin from unthreading. The final gene, *uboD*, in the BGC encodes an ABC transporter that serves as an immunity factor in the native producer *B. ubonensis* to export bioactive ubonodin outside the cell. The ubonodin structure was generated from PDB 6POR. (**B**) Cartoon summarizing essential steps in the mode of action of ubonodin. As ubonodin has an intracellular target, it must sequentially cross the outer (OM) and inner (IM) membranes of the target cell before it can bind RNA polymerase and inhibit transcription. Prior to this work, the identity of the OM transporter was not known.

We recently reported the discovery of a novel lasso peptide, ubonodin, that has potent antimicrobial activity against several *Burkholderia* pathogens (**Figure 1A**) (Cheung-Lee et al., 2020). Our work demonstrated that ubonodin inhibits RNA polymerase (RNAP) *in vitro*. Yet despite having a highly conserved cellular target, ubonodin is selectively bioactive, inhibiting some but not all *Burkholderia* strains. Because ubonodin has a cytoplasmic target, we speculated that its ability to access the interior of target cells is key to determining the extent of activity. That is, ubonodin transport across the bacterial outer membrane (OM) followed by the inner membrane (IM) is a prerequisite for RNAP engagement (**Figure 1B**). Due to its large size, cellular uptake of ubonodin likely requires membrane transporters as seen for other lasso peptides and RiPPs (Cao et al., 2021; Mathavan and Beis, 2012). Identifying the membrane transporters that are central to the mechanism of action of ubonodin will enable accurate prediction of its spectrum of activity, an important step toward developing ubonodin as an antibiotic.

Here, we first focus on the initial stage of cellular uptake, the OM transport step which is a key barrier to compound bioactivity. *Burkholderia* have unusually large, multireplicon genomes encoding multiple possible OM transporters, consistent with the ability of *Burkholderia* to survive in diverse ecological niches (Mahenthiralingam et al., 2005). The plethora of nutrient uptake systems in *Burkholderia* however complicates the search for ubonodin transporters. Overcoming this challenge, we developed a comparative genomics approach complemented with targeted mutagenesis to quickly identify the TonB-dependent transporter PupB as the ubonodin OM receptor in *B. cepacia*. For any Bcc strain, the presence of a close PupB homolog served as a reliable predictor of ubonodin susceptibility. We further show that PupB is subjected to iron-mediated transcriptional repression and regulation by a protein activator. Our finding that OM transport correlates with susceptibility provides a molecular explanation for ubonodin bioactivity. In addition to pinpointing the OM receptor, we provide evidence that ubonodin uses an ATP-powered IM transporter in the second stage of cellular uptake. Finally, beyond the focus on ubonodin, our studies reveal new insights into *Burkholderia* physiology. Our multipronged experimental and computational approach will also be useful as a general framework for other compound mechanisms of action studies.

## RESULTS

### Ubonodin inhibits the growth of select Bcc strains

In our original report on the discovery of ubonodin, we had examined its bioactivity against an initial panel of 8 *Burkholderia* strains (Cheung-Lee et al., 2020). Of these strains, only 2 – *B. cepacia* ATCC 25416 and *B. multivorans* ATCC 17616 – are members of the Bcc and both were found to be susceptible to ubonodin. Due to the limited information on bioactivity, what distinguishes ubonodin-susceptible (ubo^S^) from non-susceptible (ubo^N^) strains was not immediately clear. Thus, we decided first to more broadly examine the spectrum of ubonodin bioactivity to have a larger panel of strains for comparison. We measured the activity of ubonodin against 10 additional Bcc strains, focusing on strains from the Bcc because they are most closely related to the ubonodin producer strain, *B. ubonensis*, and RiPPs tend to have a focused spectrum of activity (Cao et al., 2021; Li and Rebuffat, 2020). Moreover, we anticipated that the elusive genetic signatures differentiating ubo^S^ from ubo^N^ Bcc strains should be more apparent as Bcc members are otherwise highly similar. In total, we tested the activity of ubonodin against 12 Bcc strains representing 6 distinct Bcc species or genomovars, including several Bcc strains that were recommended for genetic studies (Coenye et al., 2001; Depoorter et al., 2020; Mahenthiralingam et al., 2000). These Bcc species are also most associated with infections in CF patients (LiPuma, 2010; Zlosnik et al., 2020). Of the 12 Bcc strains tested, 6 were susceptible to ubonodin (**Figure 2** and **Figure 2 – figure supplement 1**). The levels of susceptibility for ubo^S^ Bcc strains were similar, ranging from 10-40 μM ubonodin (**Table 1**). Thus, ubonodin is selectively active against a subset of Bcc strains.

**Table 1.** Minimum inhibitory concentration (MIC) of ubonodin against susceptible Bcc strains. MIC measurements were obtained via M63 spot-on-lawn assay and observing for the lowest ubonodin concentration at which a zone of inhibition was clearly visible. MIC values for strains ATCC 17616 and ATCC 25416 were previously reported (Cheung-Lee et al., 2020).

| Strain | Ubonodin MIC (μM) |
| --- | --- |
| *B. multivorans* AU30438 | 10 |
| *B. multivorans* ATCC 17616 | 20 |
| *B. cepacia* ATCC 25416 | 40 |
| *B. cenocepacia* AU0756 | 10 |
| *B. cenocepacia* AU24362 | 20 |
| *B. cenocepacia* J2315 | 20 |

**Figure 2.**
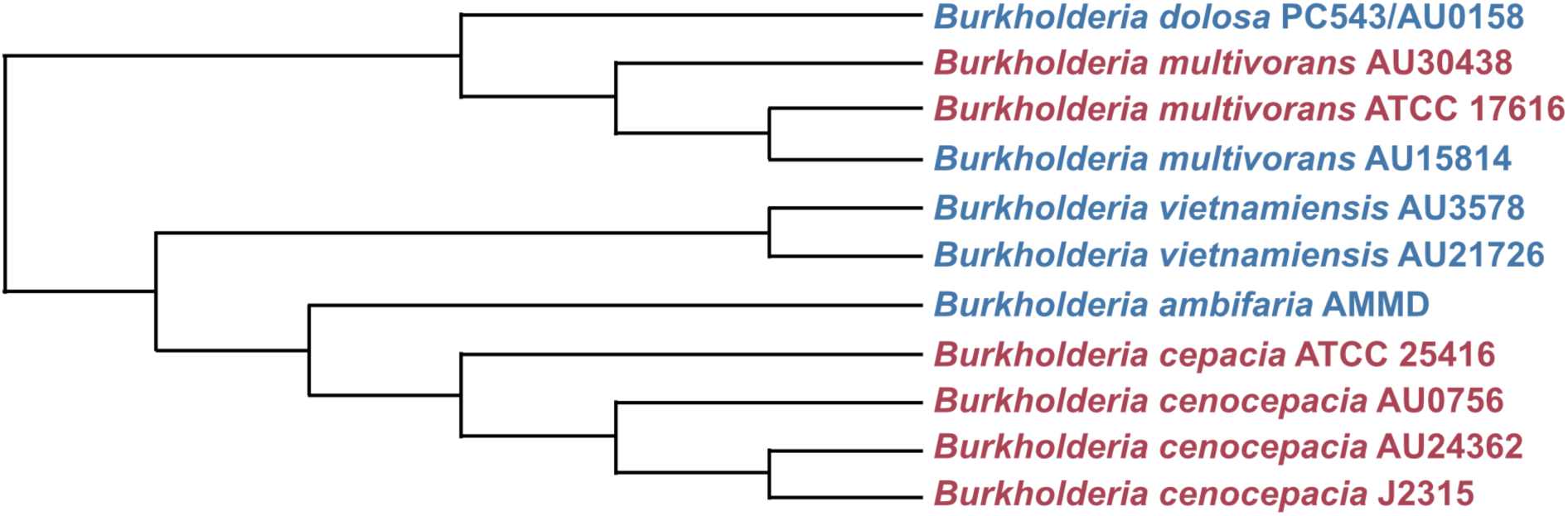
Ubonodin selectively inhibits a subset of Bcc strains. Clustal Omega alignment of 7 concatenated gene fragments (*atpD, gltB, gyrB, recA, lepA, phaC*, and *trpB*) encoded by ubonodin-susceptible (red) and non-susceptible (blue) Bcc strains was used to generate the cladogram. As Bcc strains are closely related, alignment of these housekeeping genes provides higher taxonomic resolution than 16S rRNA analysis (Depoorter et al., 2020; Vandamme and Peeters, 2014; Winsor et al., 2008). Shown is a Neighbor-joining tree without distance corrections.

**Figure 2 – figure supplement 1.**
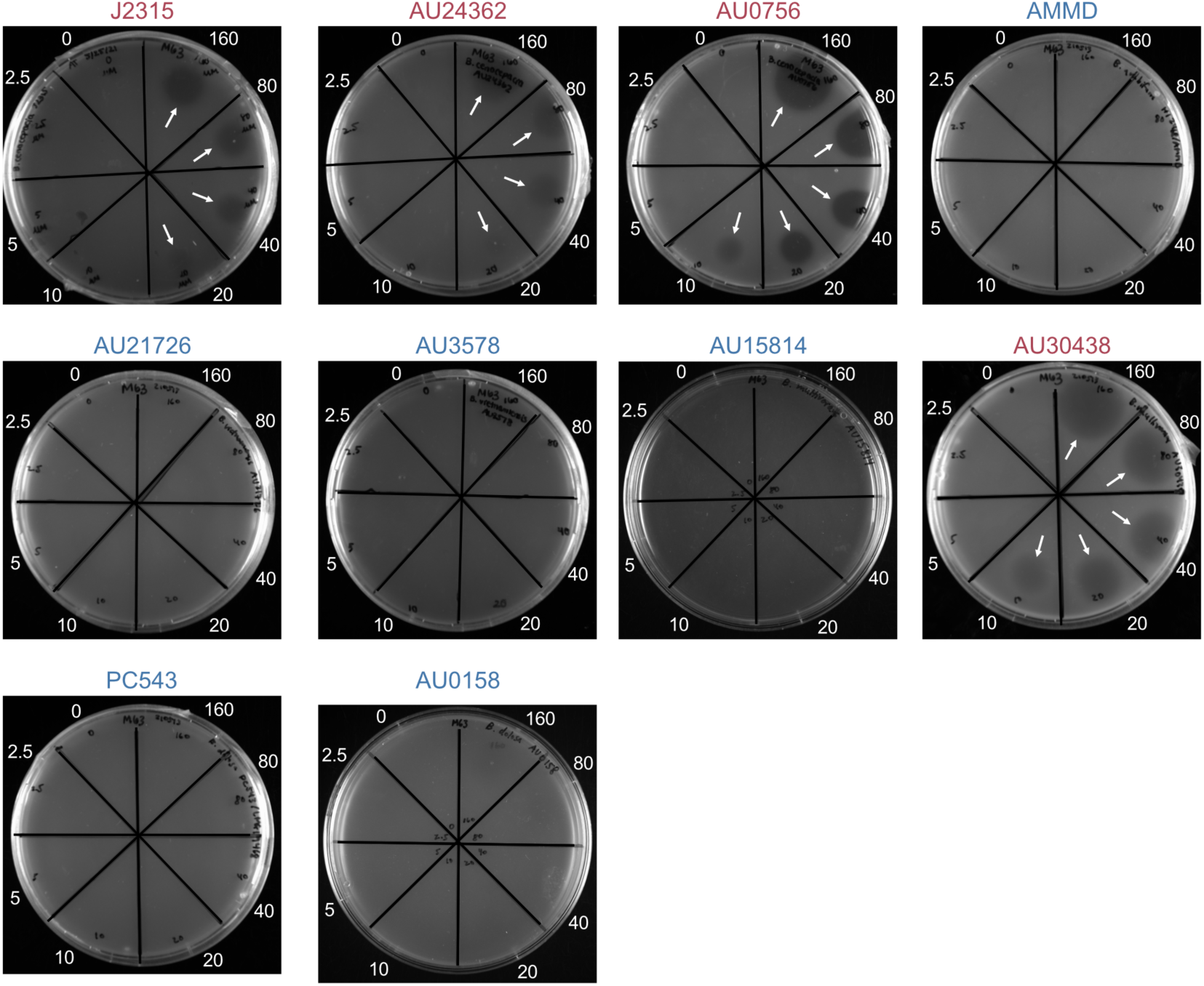
Spot-on-lawn assay assessing Bcc strains for ubonodin susceptibility. Cultures were grown to exponential phase, 10^8^ CFUs were mixed with M63 soft agar, and the mixture was plated on M63 base agar. Two-fold serial dilutions of ubonodin ranging from 0-160 μM were spotted (10 μL spots) onto the respective sectors once the cell-agar mixture had solidified. The plates were incubated at 30°C for ∼16 h until clear zones (arrows) indicating bacterial growth inhibition were observed. We note that *B. multivorans* AU15814 grew poorly on M63 compared to the other Bcc strains, which formed a lawn of cells.

We next wanted to understand how the ubo^S^ Bcc strains differ from the ubo^N^ strains. One possible explanation would be if in the ubo^N^ strains, the molecular target RNAP has specific mutations that abolish ubonodin recognition and binding. Structural work on the transcription-inhibiting lasso peptides MccJ25 (Delgado et al., 2001; Salomón and Farías, 1992; Yuzenkova et al., 2002) and capistruin (Knappe et al., 2008; Kuznedelov et al., 2011) confirmed that both peptides bind within the secondary channel of RNAP blocking the path to the catalytic center (Braffman et al., 2019). Both MccJ25 and capistruin interact with residues belonging to the RNAP β and β’ subunits. Given the structural and functional similarities between ubonodin and these other lasso peptides (Cheung-Lee et al., 2020), we reasoned that their modes of binding to RNAP would also be analogous and ubonodin would contact β and β’ subunit residues. Therefore, we compared the amino acid sequence of the β (RpoB) and β’ (RpoC) subunits for the 12 Bcc strains tested for ubonodin susceptibility to determine if ubo^N^ strains encode distinctly unique β/β’ variants from ubo^S^ strains. By multiple sequence alignment, we found that the β/β’ subunits across all Bcc strains tested were >98.0% identical (**Figure 2 – figure supplement 2**). Importantly, β/β’ variants from ubo^S^ strains are not more similar than their ubo^N^ counterparts. Ubonodin likely binds the RNAP holoenzyme of all the Bcc strains tested equivalently, meaning that other cellular properties are instead responsible for the observed difference in ubonodin susceptibility.

**Figure 2 – figure supplement 2.**
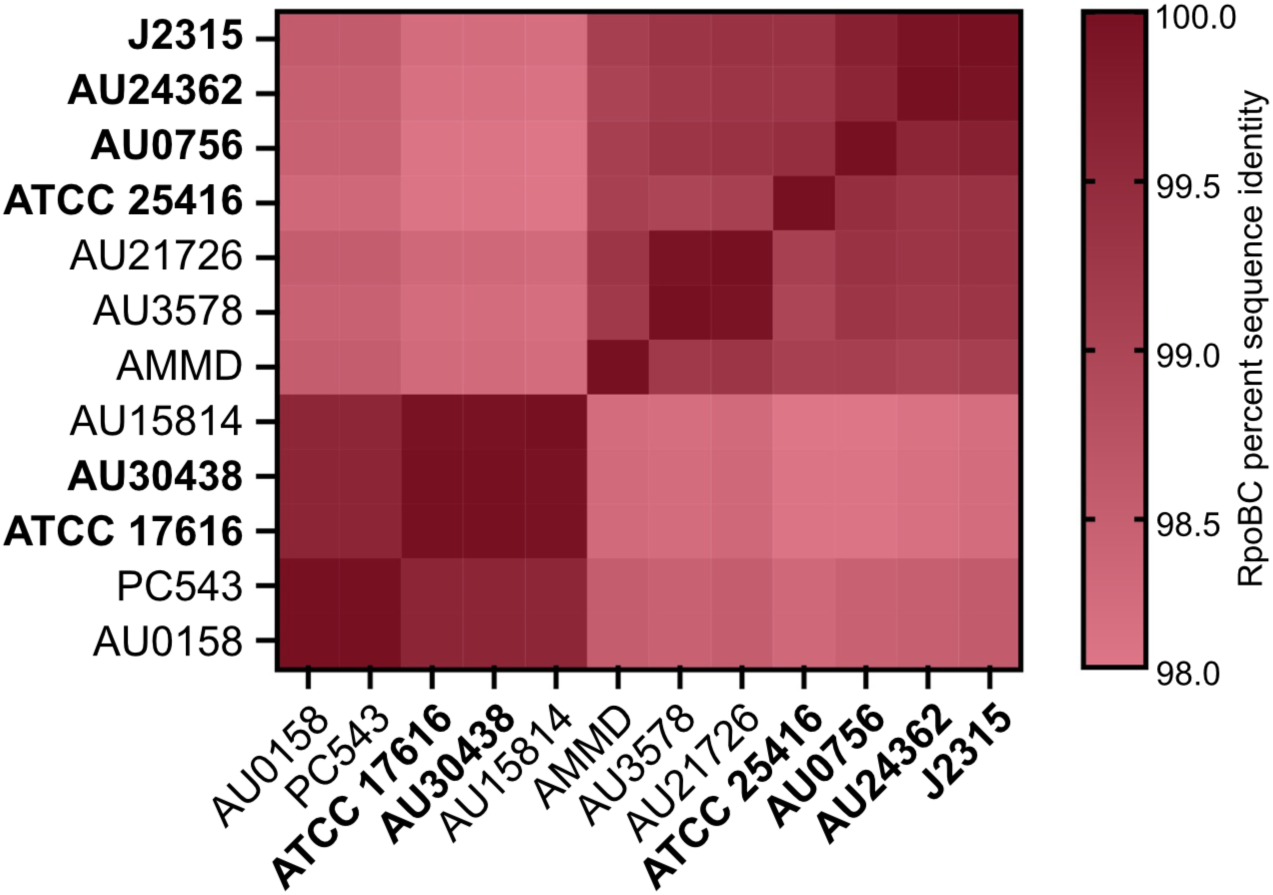
Conservation of the RNA polymerase β and β’ subunits across the Bcc strains tested for ubonodin susceptibility. Heat map showing percent sequence identity when the concatenated amino acid sequences of RpoB (β subunit) and RpoC (β’ subunit) were aligned with Clustal Omega. A darker shade represents more identical RpoBC sequences for a pair of strains along the x- and y-axes. Ubonodin-susceptible Bcc strains are bolded.

### Iron-dependent inhibition of ubonodin bioactivity reveals a specific outer membrane transport pathway for ubonodin uptake

The bioactivity of ubonodin, and indeed any compound with an intracellular target, is a product of its ability to first access and then bind its target. Previous studies have shown that cellular uptake is required for lasso peptides with RNAP-inhibiting activity to be bioactive (Cheung-Lee et al., 2019; Li et al., 2021; Metelev et al., 2017; Salomón and Farías, 1993). In all cases where the lasso peptide OM receptor has been identified, the receptors are TonB-dependent transporters (TBDTs), a ubiquitous class of bacterial OM proteins that import essential trace nutrients like iron from the environment (**Figure 3A**) (Krewulak and Vogel, 2011; Noinaj et al., 2010). The natural substrates of many TBDTs are siderophore compounds that chelate insoluble, environmental ferric iron (Fe^3+^) for passage into the cell (Hider and Kong, 2010).

**Figure 3.**
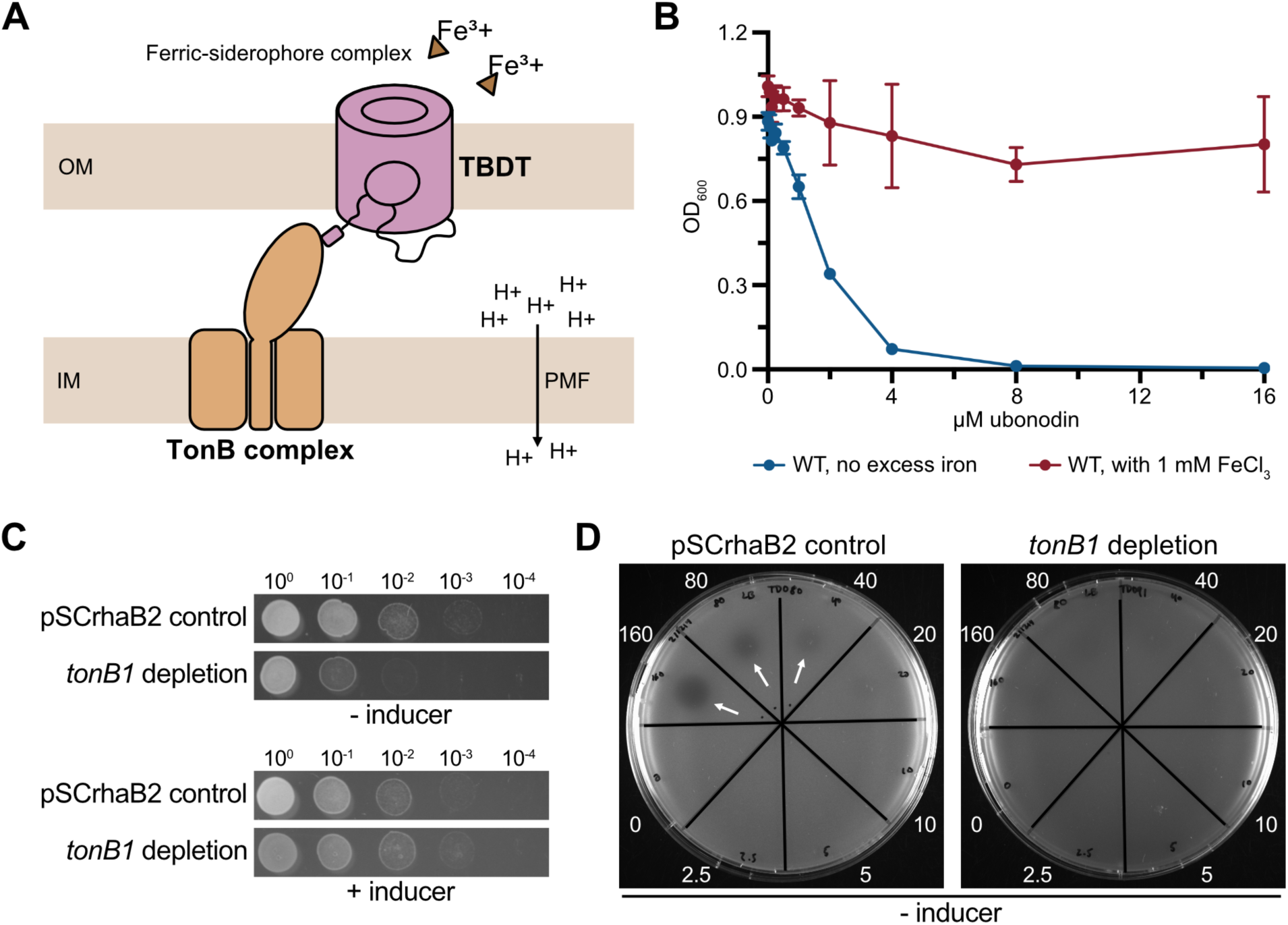
Ubonodin repurposes a specific iron transport pathway for cellular entry. (**A**) Cartoon describing a general TonB-dependent iron transport pathway. The TonB-dependent transporter (TBDT) localized to the bacterial outer membrane (OM) is activated by a partner TonB complex at the inner membrane (IM). TonB channels the energy from the proton motive force (PMF) to drive OM transport of siderophore-chelated ferric iron. (**B**) Excess iron inhibits ubonodin activity. *B. cepacia* was grown in plain LB (blue) or supplemented with excess iron (red), and the final cell density (OD_600_) was measured after ∼16 h of growth at 30°C. The mean ± standard deviation for independent biological triplicates is graphed; error bars are not apparent for points with a small standard deviation. (**C**) Depletion of *tonB1* reduces *B. cepacia* cell growth but *tonB1* is not required for viability. Cultures of *B. cepacia* propagating an empty vector (pSCrhaB2) and the *tonB1* depletion mutant were grown to exponential phase and normalized to an equivalent OD_600_. Five μL spots of ten-fold serial dilutions of the cultures were spotted onto LB agar with and without the inducer (0.2% L-rhamnose). Plates were imaged after ∼15 h of growth at 30°C. (**D**) The *B. cepacia tonB1* depletion mutant is less susceptible to ubonodin. Cultures were grown to exponential phase and 10^8^ CFUs were plated on LB agar. Ten μL of 0-160 μM of ubonodin were spotted in the respective sectors. Zones of inhibition (arrows) were observed after the spot-on-lawn plates were incubated at 30°C for ∼15 h.

To probe whether ubonodin also uses an iron receptor for passage across the *B. cepacia* OM, we examined how excess iron affects ubonodin activity. Bacterial iron transport systems are highly regulated to prevent toxic accumulation of intracellular iron which can lead to oxidative damage (Andrews et al., 2003; Noinaj et al., 2010). In iron-depleted environments, TBDT and siderophore biosynthesis genes are upregulated to maximize the ability to scavenge iron; conversely, these genes are downregulated in iron-rich environments (Butt and Thomas, 2017; Noinaj et al., 2010; Thomas, 2007). If ubonodin enters *B. cepacia* through iron-regulated TBDTs, we expected that excess iron would antagonize ubonodin activity by reducing transporter abundance. When *B. cepacia* was grown in media supplemented with excess iron, ubonodin indeed could no longer robustly inhibit cell growth, suggesting that ubonodin uptake occurs through one or more iron-regulated OM transporters (**Figure 3B**).

We next probed the specific role of TBDTs in ubonodin transport. To our knowledge, a comprehensive list of *B. cepacia* TBDTs is not available. Thus, we first performed protein BLAST search using *Escherichia coli* FhuA, the native receptor for MccJ25 (Salomón and Farías, 1993), and 34 known and predicted TBDTs from *Pseudomonas aeruginosa* (Luscher et al., 2018) to identify *B. cepacia* TBDT candidates. The analysis revealed 29 TBDT homologs in *B. cepacia* strain ATCC 25416 ranging from ∼13% to ∼37% sequence identity to *E. coli* FhuA (**Supplementary file 1 – Supplementary Table 1**). Any number or none of these predicted TBDTs might be involved in ubonodin import, but all should need TonB to function. Identifying and knocking out the TonB regulator would avoid redundancy of the TBDTs when probing their role in ubonodin transport.

As the identity of TonB in *B. cepacia* is also unknown, we first searched for homologs to known TonB sequences from *B. cenocepacia* strain K56-2 (Pradenas et al., 2017) and *B. mallei* strain ATCC 23344 (Mott et al., 2015). Protein BLAST search yielded 4 hits with the top hit encoded by the *B. cepacia* gene GGFLHMPP_02544 (*tonB1*) (**Supplementary file 1 – Supplementary Table 2**). When we attempted to knock out *tonB1* using one-step allelic replacement (Shastri et al., 2017), we could only isolate single-crossover mutants that still retained the wild-type allele, likely because TonB is important for *Burkholderia* cellular fitness (Higgins et al., 2017; Mott et al., 2015; Pradenas et al., 2017). Instead, we constructed a *B. cepacia tonB1* depletion mutant in which the chromosomal copy of *tonB1* is deleted while rhamnose-inducible extrachromosomal copies of *tonB1* are provided on a replicative plasmid (Cardona and Valvano, 2005). As expected, when the inducer was withheld to deplete *tonB1*, the mutant was viable but grew poorly compared to *B. cepacia* WT (**Figure 3C**). Growth was restored to near-WT levels either in the presence of L-rhamnose or when the cultures were supplemented with ferrous iron (Fe^2+^) to counteract iron starvation due to the defect in iron acquisition, as previously demonstrated (Mott et al., 2015; Pradenas et al., 2017) (**Figure 3 – figure supplement 1**). With this mutant in hand, we found that *tonB1* depletion led to ubonodin resistance (**Figure 3D and Figure 3 – figure supplement 2**). By contrast, loss of the other 3 TonB homologs, which are non-essential, did not impact ubonodin susceptibility because the Δ*tonB2*, Δ*tonB3*, and Δ*tonB4* mutants were equally susceptible to ubonodin as WT (**Table 2 and Figure 3 – figure supplement 3**). Taken together, these findings show that ubonodin uses one specific TonB-dependent pathway to cross the *B. cepacia* OM.

**Table 2.** Minimum inhibitory concentration of ubonodin for *B. cepacia tonB* mutants. MIC was defined as the lowest concentration of ubonodin required to inhibit bacterial growth via spot-on-lawn assay. The *tonB1* depletion mutant was compared to the pSCrhaB2 empty vector control strain on LB agar ± the inducer (0.2% L-rhamnose). For the *tonB2, tonB3*, and *tonB4* mutants for which the gene could be deleted, MICs were assessed against *B. cepacia* WT on plain LB agar. na indicates not applicable.

| MIC of ubonodin (μM) |  |  |
| --- | --- | --- |
| Strain | - inducer | + inducer |
| pSCrhaB2 control | 40 | 40 |
| *tonB1* depletion | >160 | 40 |
| WT | 20 | na |
| $\Delta tonB2$ | 20 | na |
| $\Delta tonB3$ | 20 | na |
| $\Delta tonB4$ | 20 | na |

**Figure 3 – figure supplement 1.**
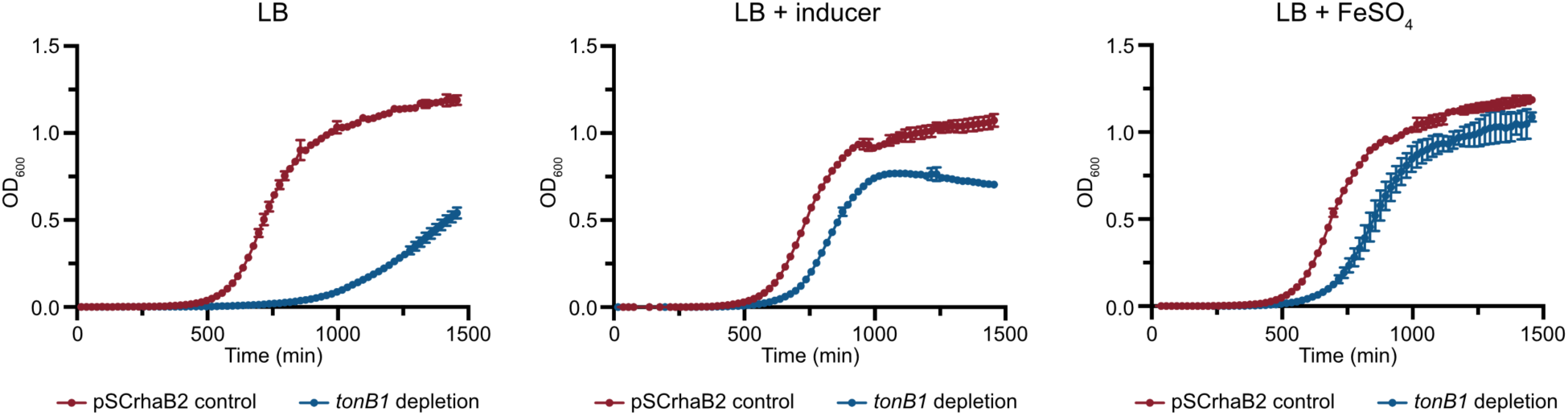
Iron supplementation mitigates the defect in *B. cepacia* cell growth due to *tonB1* depletion. The *tonB1* depletion mutant (blue) and empty vector control strain (red) were grown in LB (left graph) and in the presence of 0.02% L-rhamnose (middle graph) or 200 μM FeSO_4_ (right graph). The mean ± standard deviation for technical triplicates from the same overnight culture is shown.

**Figure 3 – figure supplement 2.**
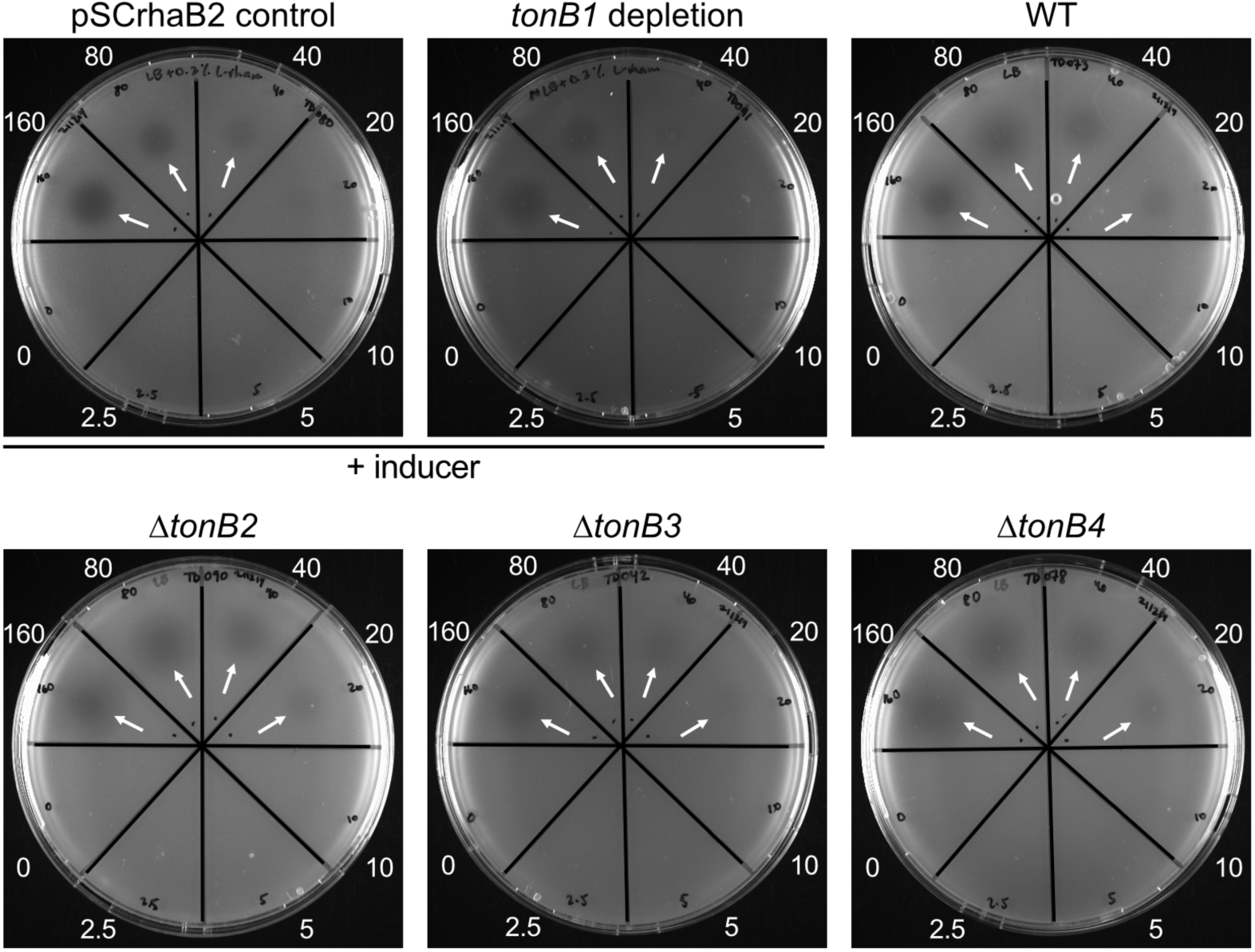
A single TonB pathway is required for ubonodin activity. Spot-on-lawn LB plates with 10^8^ CFUs of the *tonB1* depletion and empty vector control strains, *B. cepacia* WT, and the deletion mutants of the other 3 *tonB* homologs were examined for susceptibility to 0-160 μM of ubonodin. Ubonodin inhibition was observed (arrows) when *tonB1* was expressed (+ 0.2% L-rhamnose) and loss of the other TonBs did not impact the MIC.

**Figure 3 – figure supplement 3.**
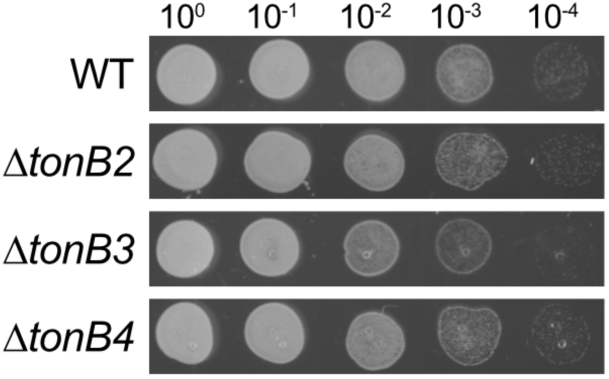
Most TonB homologs are non-essential in *B. cepacia*. Spot dilution assay showing 10-fold serial dilutions of exponential-phase cultures. Five μL of each dilution was spotted onto LB agar and the plates were incubated for ∼15 h at 30°C.

### Comparative genomics identifies siderophore receptors unique to ubonodin-susceptible Bcc strains

The next challenge was to identify the exact TBDT(s) that TonB1 presumably activates to enable ubonodin transport. All Bcc strains encode multiple TBDTs but the set of TBDTs found in each strain is variable (Butt and Thomas, 2017). Reasoning that ubo^S^ Bcc strains encode a distinct set of TBDTs from ubo^N^ Bcc strains, we developed a comparative genomics approach to predict the TBDTs that are unique to ubo^S^ strains (**Figure 4A**). These TBDTs should include any ubonodin transporter(s) and their absence in ubo^N^ strains would explain why those strains are naturally non-susceptible. Our expanded panel of Bcc strains tested for ubonodin susceptibility provided a dataset sufficient for comparison. First, we built a protein BLAST database for each Bcc strain and queried the 29 predicted *B. cepacia* ATCC 25416 TBDTs against each database. For each TBDT, we calculated the percent similarity normalized by query coverage between the TBDT and its highest-scoring alignment across all the Bcc strains tested. We then manually examined the heatmap summarizing the BLAST scores for *B. cepacia* TBDTs that show high sequence conservation only or predominantly in the other ubo^S^ strains.

**Figure 4.**
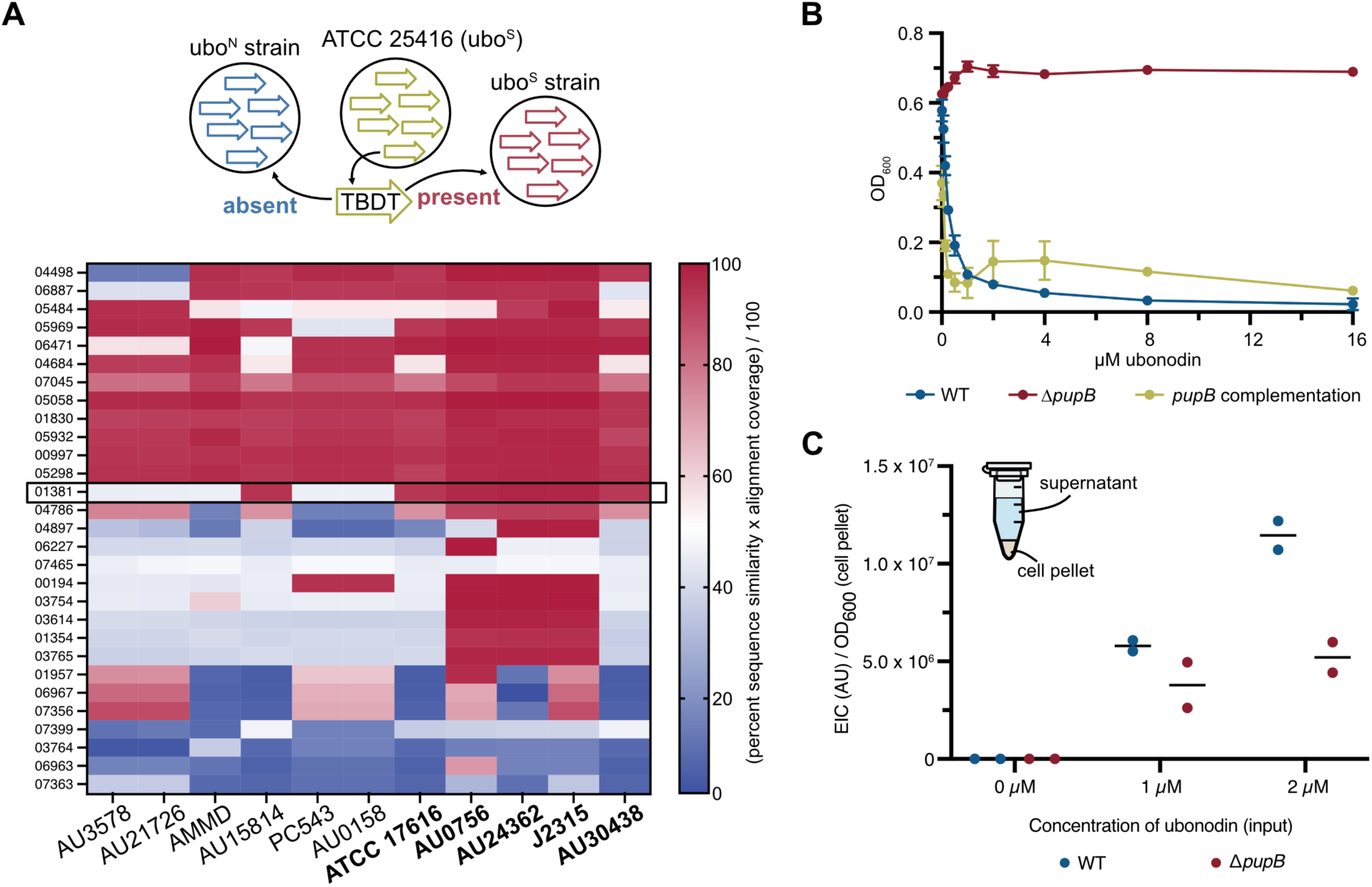
Phenotype-guided comparative genomics identifies PupB as the *Burkholderia* outer membrane receptor for ubonodin. (**A**) Comparative genomics revealed TBDT genes which are conserved in subsets of Bcc strains tested for ubonodin susceptibility. Top: Our comparative genomics approach used protein BLAST to search for the closest homolog of each putative ATCC 25416 TBDT in each ubonodin-susceptible (ubo^S^) and non-susceptible (ubo^N^) Bcc strain. Bottom: For each TBDT search, the percent sequence similarity of the top hit normalized to the alignment coverage was calculated. Each row of the heat map represents one of the 29 TBDTs assessed and each column represents one of the 11 Bcc strains besides the *B. cepacia* seed strain tested for ubonodin susceptibility. The blue-to-red spectrum indicates low-to-high protein sequence conservation. Only one hit encoded by the gene GGFLHMPP_01381 was predominately conserved in ubo^S^ (bolded) but not ubo^N^ (non-bolded) strains. (**B**) PupB is required for ubonodin activity. Exponential-phase cultures were diluted to a starting OD_600_ of 0.0005 in cation-adjusted Mueller-Hinton II broth and exposed to 0-16 μM of ubonodin. Endpoint OD_600_ was measured after ∼16 h of growth at 30°C. Complementation of *pupB* was achieved by providing *pupB* on the pSCrhaB2 plasmid with 0.0002% L-rhamnose induction. Plots show the mean ± standard deviation for independent biological triplicates; error bars are not apparent for points with a small standard deviation. (**C**) Loss of PupB reduces ubonodin uptake. *B. cepacia* WT and Δ*pupB* endpoint cultures assessed for inhibition with sub-MIC concentrations of ubonodin were also analyzed for ubonodin uptake. The endpoint cultures were centrifuged to separate the supernatant and cell pellet, and cell lysates prepared by cold methanol extraction were analyzed by LC-MS analysis to measure the amount of ubonodin internalized by the cells. The sum of the extracted ion count (EIC) for various ubonodin species observed by LC-MS was normalized to the final cell density. Non-specific absorption to the cell surface might account for some background ubonodin signal. Two biological replicates were independently measured.

### The *B. cepacia* siderophore receptor PupB is the ubonodin outer membrane transporter

One *B. cepacia* TBDT homolog encoded by the gene GGFLHMPP_01381 (*pupB*) stood out because it was predicted to be highly conserved in all ubo^S^ strains and only one ubo^N^ strain (**Figure 4A**). PupB is named for its resemblance to TBDTs that transport pseudobactin-type siderophores (Koster et al., 1993). Overall, PupB satisfies the conservation pattern expected for an ubonodin transporter that would define the spectrum of activity. No other TBDT homolog showed a similar conservation pattern. To test if PupB is involved in ubonodin transport, we deleted *pupB* in *B. cepacia* and found that the mutant was completely resistant to ubonodin, as expected with loss of cellular uptake (**Figure 4B**). Ubonodin susceptibility was restored when *pupB* was provided back in *trans* on a plasmid. These results strongly implicate PupB in ubonodin OM import. We also carried out cellular uptake assays comparing *B. cepacia* WT to the Δ*pupB* mutant. *B. cepacia* was incubated with ubonodin and intracellular levels of ubonodin were measured by LC-MS analysis of the cell lysates, normalized to the endpoint cell density for each sample. In summary, we found that ubonodin accumulated to higher levels in WT cells and cellular uptake of ubonodin was reduced in the absence of PupB (**Figure 4C**). Our genetic and biochemical results conclusively demonstrate that ubonodin uses PupB as an OM receptor for initial cellular entry.

We also examined the PupB sequences in more detail to understand how the homologs belonging to the ubo^S^ strains are unique. Alignment of the closest PupB homologs for all 12 Bcc strains tested revealed a distinct N-terminal motif found almost exclusively in the ubo^S^ strains (**Figure 4 – figure supplement 1**). In addition to the β-barrel domain and the periplasmic plug domain that sterically blocks the β-barrel lumen, a subset of TBDTs harbor an additional domain at the N-terminus (Noinaj et al., 2010). This N-terminal extension is involved in signaling through a cognate IM σ regulator-extracytoplasmic σ factor pair to regulate the expression of the parent TBDT and related transport genes (Braun et al., 2003; Braun and Mahren, 2005; Visca et al., 2002). As intracellular iron concentrations must be precisely controlled, this added level of regulation provides a useful feedback mechanism. Besides the N-terminal extension, the PupB hits were otherwise highly similar across all strains. That PupB homologs harbor an additional N-terminal signaling domain is especially interesting in the context of the genomic location of the *pupB* gene, which we will further discuss below.

**Figure 4 – figure supplement 1.**
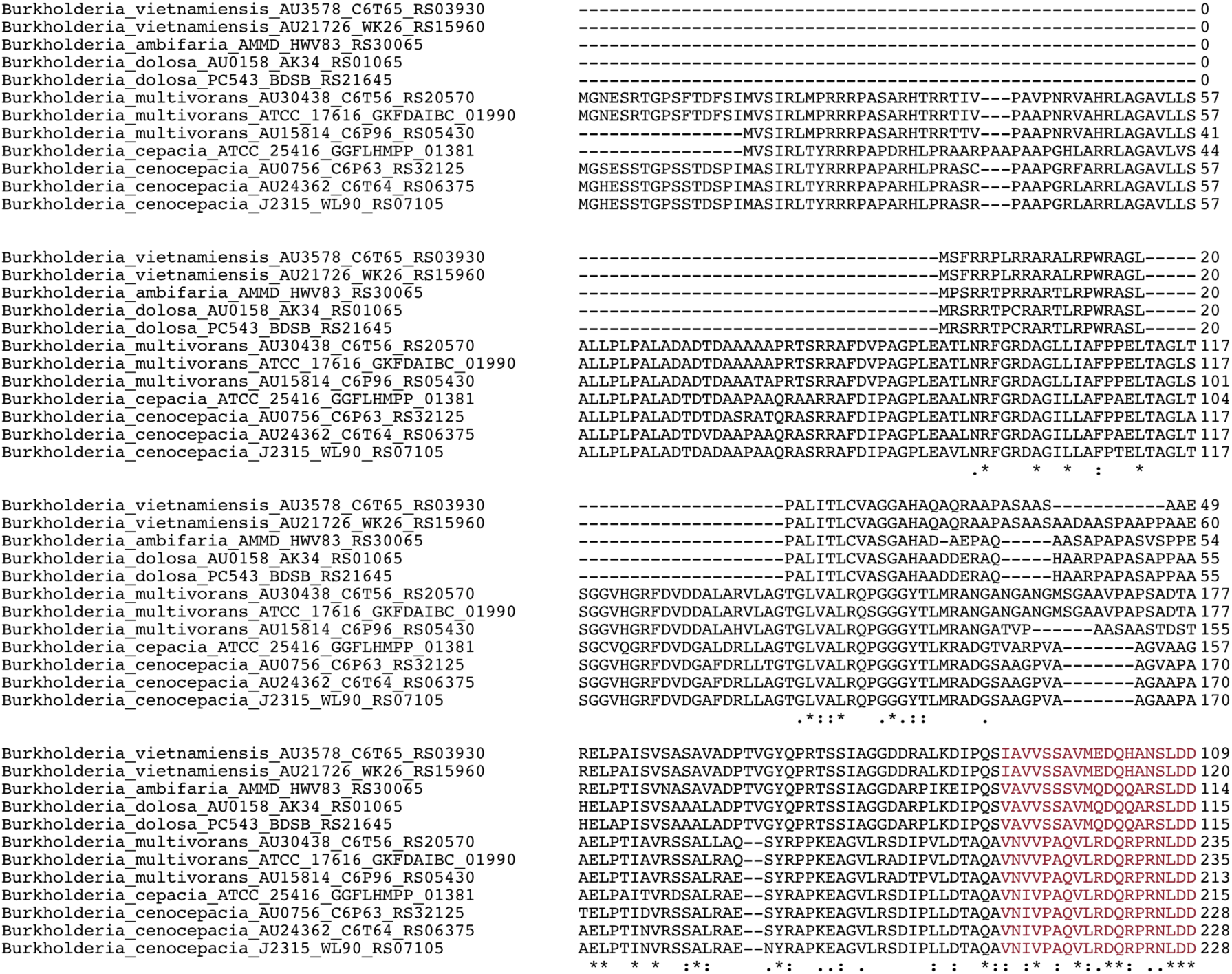
PupB homologs encoded by ubonodin-susceptible strains harbor an N-terminal extension domain. The closest PupB homologs found by protein BLAST search against the 12 Bcc strains tested for ubonodin susceptibility were aligned with Clustal Omega. Only the N-terminal portion of the alignment is shown here. The start of the plug domain based on protein BLAST annotation of the ATCC 25416 PupB homolog is denoted in red text.

### RNA-sequencing of *B. cepacia* profiles the global cellular response to excess iron

Having identified the ubonodin OM receptor, we revisited the observation that excess iron inhibits ubonodin activity to understand if excess iron does so by repressing *pupB* expression, thereby impeding cellular uptake. For this purpose, we performed RNA-seq to compare the *B. cepacia* transcriptome between 2 growth conditions: standard LB with baseline iron and supplemented with 1 mM FeCl_3_ to match the conditions of the ubonodin inhibition assay. An advantage of our RNA-seq approach is that even if excess iron does not directly impact *pupB* expression, we might still understand how PupB transport could be affected via possible changes in the expression of other transport pathway components, *e*.*g*., the TonB regulator.

As expected, growth in excess iron led to the repression of 183 genes, among them iron transport and utilization genes and siderophore biosynthesis genes (**Figure 5A and Supplementary file 1 – Supplementary Table 3**). Several of these iron pathway-related genes were accordingly found to be induced under low-iron growth conditions in prior transcriptomics studies (Sass et al., 2013; Tuanyok et al., 2005; Tyrrell et al., 2015). By contrast, 114 genes were significantly upregulated in the presence of excess iron, with none annotated as encoding for iron transporters (**Figure 5A and Supplementary file 1 – Supplementary Table 4**). Consistent with the observed gene expression changes, we noticed that *B. cepacia* cultures grown in excess iron were paler than the yellow-green cultures grown in standard rich media (**Figure 5 – figure supplement 1**). As the yellow-green color is a property of fluorescent iron-scavenging siderophores (Thomas, 2007), the color change supports that siderophore production was downregulated in response to excess iron.

**Figure 5.**
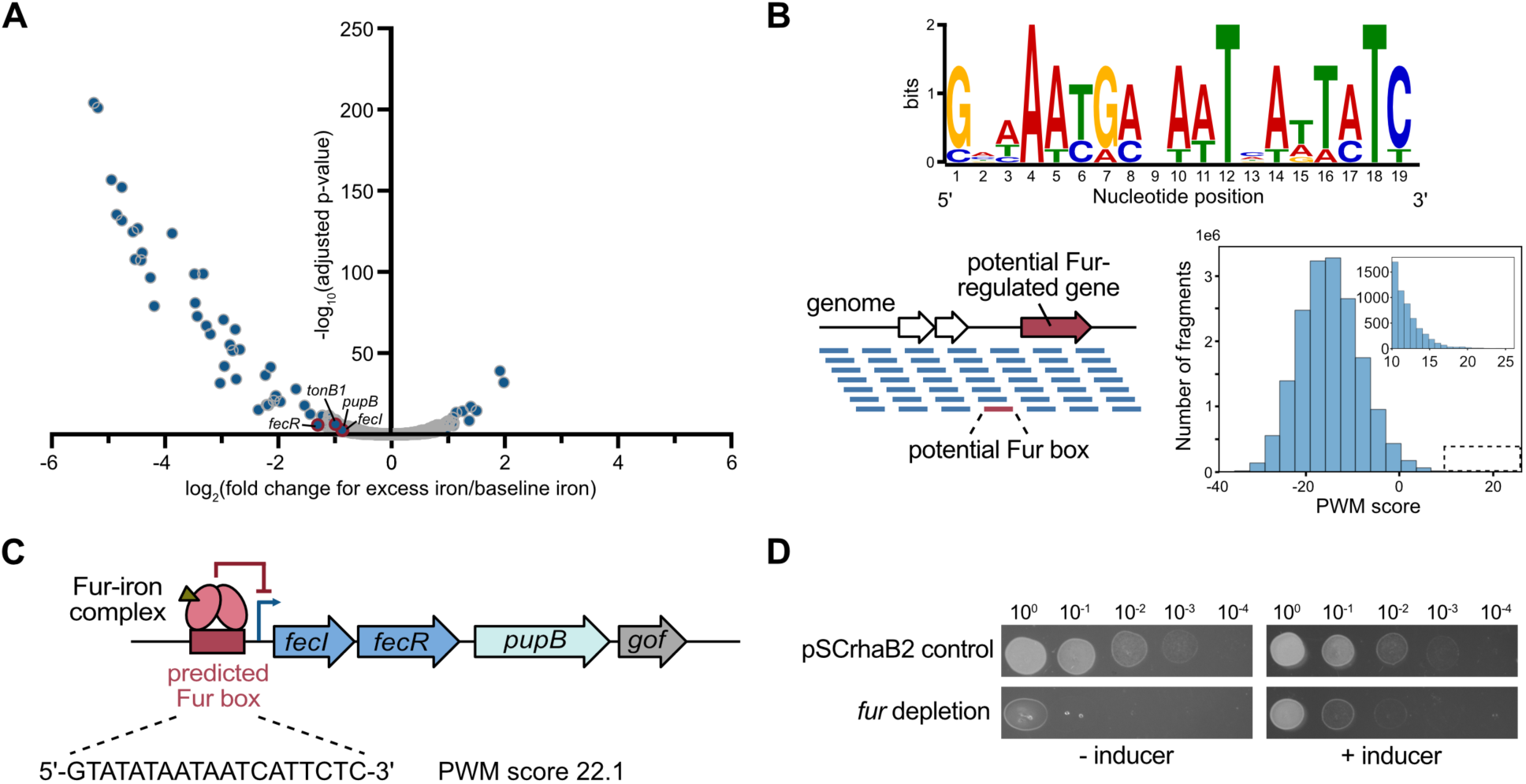
PupB is a member of the *Burkholderia* iron regulon. (**A**) Volcano plot showing iron-repressed and upregulated genes in *B. cepacia* ATCC 25416. Total RNA was extracted from independent biological triplicate cultures of *B. cepacia* WT grown to mid-exponential phase in LB ± 1 mM FeCl_3_. Illumina RNA-seq libraries were prepared and sequenced. Genes which exhibited significant changes (adjusted p-value < 0.01) in read count between the 2 growth conditions are shown as blue dots superimposed onto all detectable genes in gray. Select genes of interest in the PupB transport pathway are denoted with a red outline and labeled. (**B**) Prediction of the *B. cepacia* Fur regulon. Top: Seven experimentally determined *P. aeruginosa* Fur DNA-binding sites (Dudek and Jahn, 2021; Hassett et al., 1997; Ochsner et al., 2000, 1995) were aligned using the MEME (Multiple EM for Motif Elicitation) tool to generate the Fur consensus sequence (Bailey et al., 2009). Bottom left: A position-weight matrix (PWM) built using the Fur DNA-binding sites was used to scan the ATCC 25416 genome for similar sequences indicating a potential Fur box. Bottom right: Histogram of PWM scores for all scanned genome fragments with the inset showing the distribution of scores >10. (**C**) A potential Fur box is present upstream of the 4-gene operon predicted to encode *pupB*. The sequence and PWM score of the identified Fur box are denoted. The Operon-mapper web server was used for operon prediction (Taboada et al., 2018). The last gene in the predicted operon is a gene of unknown function (*gof*). (**D**) Fur is essential in *B. cepacia*. Five-μL spots of exponential-phase cultures normalized to the same starting OD_600_ and serially diluted ten-fold were spotted on LB agar ± 0.2% L-rhamnose. Plates were incubated at 30°C for ∼15 h.

**Figure 5 – figure supplement 1.**
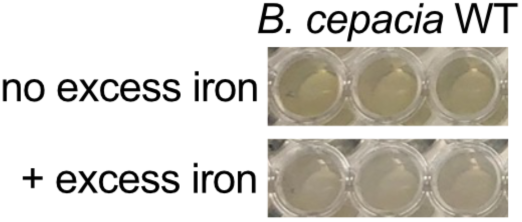
Excess iron triggers a change in the color of *B. cepacia* cultures. Overnight cultures of *B. cepacia* WT were grown in LB with (bottom) or without (top) 1 mM FeCl_3_. Whereas WT cultures are normally green, cultures exposed to excess iron were pale in color.

We examined the list of iron-repressed genes in more detail to identify any overlap with known components of the ubonodin transport pathway. Strikingly, both *pupB* (alternate gene designation, APZ15_10615) and *tonB1* (APZ15_16310) were significantly repressed when *B. cepacia* was grown in the presence of excess iron (**Figure 5A and Supplementary file 1 – Supplementary Table 3**). Of note, iron-mediated gene repression was not a general attribute for all the TBDT genes because only 8 of the 29 predicted *B. cepacia* TBDT genes showed any significant expression changes (**Supplementary file 1 – Supplementary Table 5**). The most highly repressed TBDT gene is associated with the biosynthetic operon of ornibactin, a major siderophore produced by *Burkholderia* (Darling et al., 1998; Sokol et al., 2000; Stephan et al., 1993; Thomas, 2007). That both PupB and its TonB activator are downregulated likely explains why excess iron inhibits ubonodin activity. Excess iron triggers gene expression changes that reduce the abundance of the ubonodin OM transport machinery, in turn likely reducing cellular uptake of the peptide and therefore activity.

### A strong Fur DNA-binding site is predicted upstream of the *pupB* operon

Our finding that excess iron negatively regulates *pupB* and *tonB1* expression points to an underlying iron-responsive signaling pathway in *B. cepacia*. As previously discussed, bacteria have sophisticated mechanisms to maintain iron homeostasis in cells. Central to bacterial iron homeostasis is a highly conserved transcription factor called the ferric <u>u</u>ptake regulator (Fur) protein (Escolar et al., 1999; Schäffer et al., 1985). Fur represses the transcription of genes within its regulon when bound to Fe^2+^, but in the absence of the Fe^2+^ cofactor when iron is depleted, Fur regulon genes are derepressed (Bagg and Neilands, 1987). Likewise, excess iron might repress *pupB* and *tonB* expression through Fur, but overall, there is limited information on Fur-regulated genes in *Burkholderia* (Sass and Coenye, 2020; Yuhara et al., 2008).

As an initial approach to determine if *pupB* and/or *tonB1* gene expression is Fur-regulated, we applied bioinformatics to predict the Fur regulon in *B. cepacia*. We leveraged the fact that the 19-bp Fur binding site sequence (*i*.*e*., Fur box) (Lorenzo et al., 1987) is conserved and well-established to scan the *B. cepacia* ATCC 25416 genome for candidate Fur-regulated genes (**Figure 5B and Supplementary file 1 – Supplementary Table 6**). From this analysis, a high-scoring sequence that closely matches the Fur consensus sequence was identified immediately upstream of a predicted 4-gene operon (Taboada et al., 2018) harboring *pupB*, suggesting that *pupB* is a member of the Fur regulon (**Figure 5C and Supplementary file 2**). As expected with co-regulated genes, excess iron also repressed the expression of the two genes residing upstream of *pupB* in the same operon, *fecI* (APZ15_10605) and *fecR* (APZ15_10610) (**Figure 5A and Supplementary file 3**). In contrast, no probable Fur box was predicted directly upstream of *tonB1*.

Because the ubonodin OM receptor PupB appears to be transcriptionally regulated by the Fur repressor, we decided to delete *fur* (GGFLHMPP_00610) and test if the mutant becomes hyper-susceptible to ubonodin. Our attempts to delete *fur* in *B. cepacia* were unsuccessful via one-step allelic replacement but we were able to construct a *fur* depletion mutant. When the inducer was withheld, growth of the *fur* depletion mutant was severely attenuated and only partially restored when the inducer was reintroduced, showing that Fur is essential in *B. cepacia* (Figure 5D). As the mutant was already very sick, we could not clearly determine if it is hypersensitive to ubonodin. Additional studies are needed to pinpoint the role of Fur in regulating *pupB* expression.

### Competence for outer membrane transport is key to ubonodin bioactivity

So far, we have shown that PupB is necessary for ubonodin bioactivity, but we wondered if PupB-mediated transport is also sufficient to confer ubonodin susceptibility provided that RNAP is conserved. We previously reported that while ubonodin inhibits *E. coli* RNAP activity *in vitro*, it was not bioactive against *E. coli* (Cheung-Lee et al., 2020). Speculating that the lack of bioactivity was due to exclusion of ubonodin from *E. coli* cells, we hypothesized that we could sensitize *E. coli* to ubonodin by heterologously expressing *B. cepacia* PupB together with its TonB regulator. Indeed, when PupB was overexpressed in *E. coli*, the once non-susceptible strain became susceptible to ubonodin at concentrations as low as 80 μM, which is comparable to the levels of susceptibility observed for *Burkholderia* strains (**Figure 6**). Interestingly, *B. cepacia* TonB1 was not needed to achieve susceptibility, as expression of PupB alone made *E. coli* susceptible to ubonodin and co-expression with TonB1 did not further reduce the MIC (**Figure 6 – figure supplement 1**). OM transport is therefore a key determinant of bacterial susceptibility to ubonodin and PupB clearly plays a central role in transport which cannot be readily substituted by other transporters.

**Figure 6.**
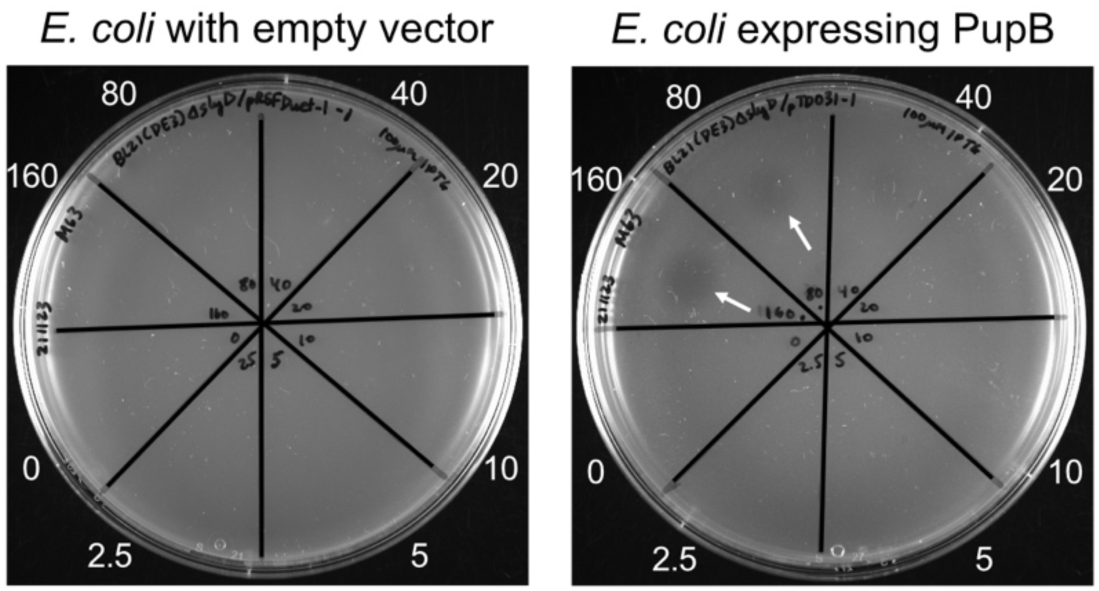
Outer membrane transport determines ubonodin susceptibility. *E. coli* BL21(DE3) Δ*slyD* propagating an empty expression plasmid or a plasmid encoding *B. cepacia pupB* was grown to exponential phase. 10^8^ CFUs were mixed with M63 soft agar containing 100 μM IPTG to induce protein overexpression and plated on M63 base agar. Ten μL of 0-160 μM ubonodin were spotted and zones of growth inhibition (arrows) were observed after ∼15 h incubation at 30°C.

**Figure 6 – figure supplement 1.**
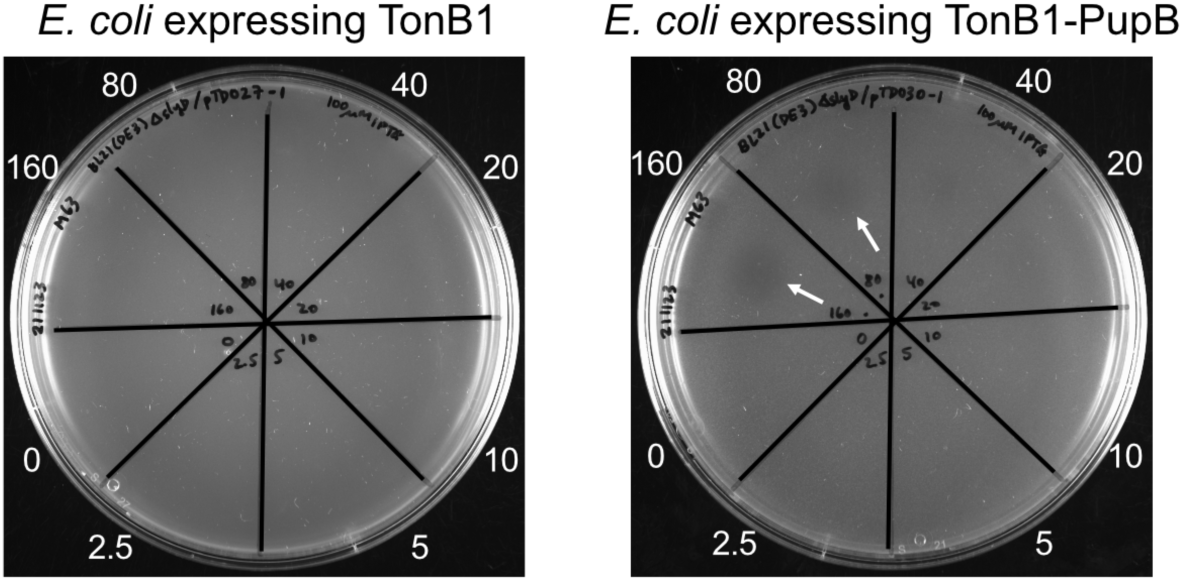
Expression of PupB alone is sufficient to confer ubonodin susceptibility. Spot-on-lawn M63 plates of 10^8^ CFUs of *E. coli* BL21(DE3) Δ*slyD* expressing *B. cepacia* TonB1 (left) together with PupB (right). Protein overexpression was achieved by the addition of 100 μM IPTG to the soft agar. Ten μL of 0-160 μM ubonodin were spotted and growth inhibition (arrows) was observed after ∼15 h incubation at 30°C.

### The putative ABC transporter YddA is the ubonodin inner membrane transporter

The ability to sensitize otherwise resistant *E. coli* to ubonodin by enabling OM uptake without interfering with IM uptake suggests that there is an *E. coli* IM transporter that can substitute for the one found natively in *B. cepacia*. The *E. coli* polytopic IM protein SbmA is known to be required for cellular uptake of the lasso peptides microcin J25 and citrocin (Cheung-Lee et al., 2019; Salomón and Farías, 1995). Thus, we wondered if there exists a homolog of SbmA in *B. cepacia* and whether such a homolog is involved in ubonodin transport. Through protein BLAST search, we found that the closest SbmA homolog in *B. cepacia* ATCC 25416 is a predicted ATP-binding cassette (ABC) transporter encoded by the gene GGFLHMPP_00373 (*yddA*). While the YddA protein sequence is only ∼25% identical to that of *E. coli* SbmA and YddA contains a cytoplasmic nucleotide binding domain not present in SbmA, YddA is highly conserved (>80% sequence identity) across all the Bcc strains tested for ubonodin susceptibility (**Figure 7A**). Consistent with YddA acting as an ubonodin IM transporter, the PSORTb bacterial protein subcellular localization prediction tool (Yu et al., 2010) localized YddA to the IM and the Phobius topology prediction tool (Käll et al., 2007, 2004) predicted 6 transmembrane helices. When we deleted *yddA* in *B. cepacia*, the bacterium was no longer susceptible to ubonodin like how *E. coli* SbmA loss-of-function mutants are resistant to microcin J25 and citrocin (**Figure 7B**) (Cheung-Lee et al., 2019; Salomón and Farías, 1995). These results strongly implicate *B. cepacia* YddA in ubonodin IM transport.

**Figure 7.**
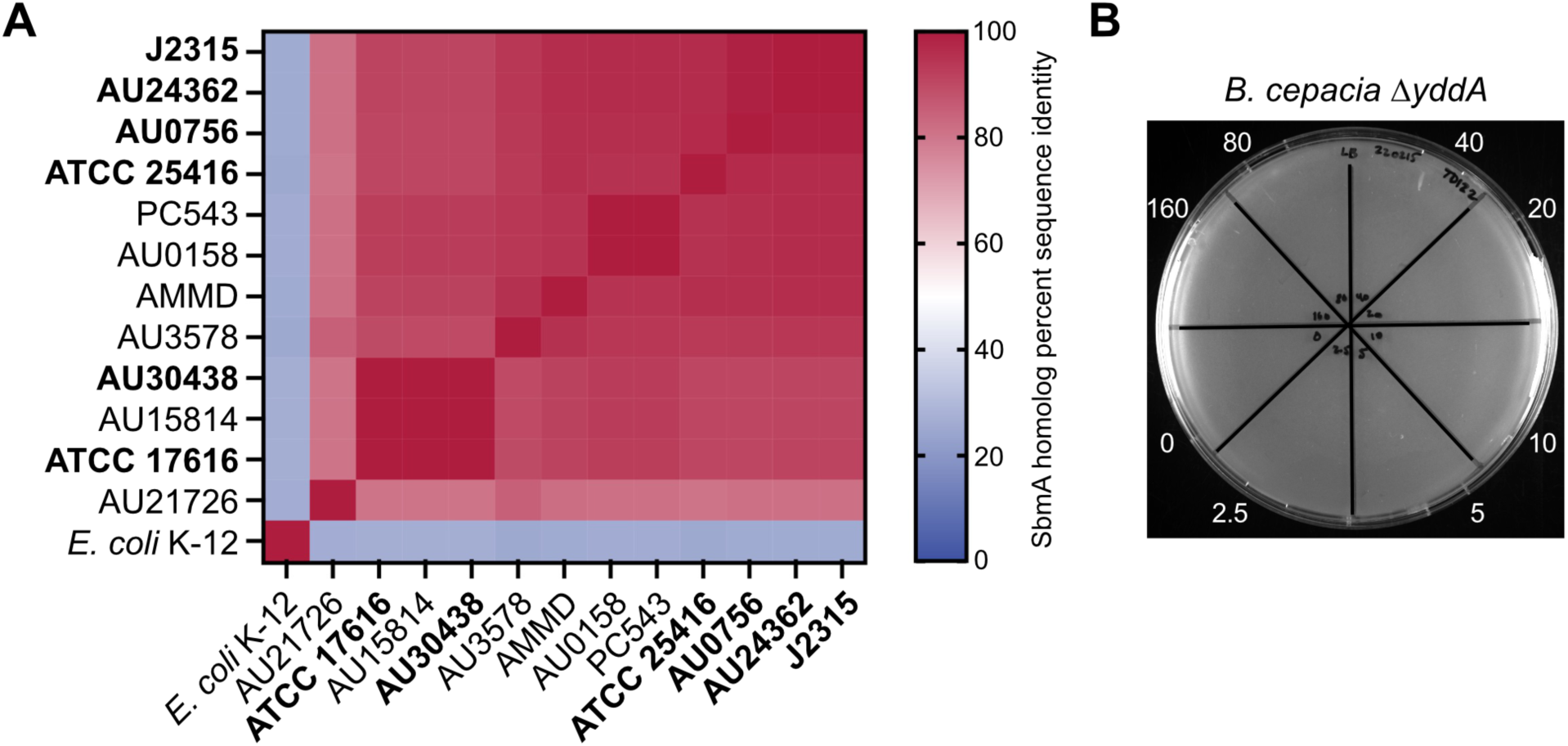
Ubonodin requires the *B. cepacia* SbmA homolog YddA for bioactivity. (**A**) Conservation of the inner membrane protein SbmA across the Bcc strains tested for ubonodin susceptibility. First, the closest homolog in strain ATCC 25416 to *E. coli* SbmA was identified using protein BLAST. This homolog, *B. cepacia* YddA, was then used to identify the closest homolog in each of the other Bcc strains. Heat map shows percent sequence identity when the SbmA homologs were aligned with Clustal Omega. A darker shade represents more identical SbmA sequences for a pair of strains along the x-and y-axes. Ubonodin-susceptible strains are bolded. (**B**) Loss of YddA makes *B. cepacia* resistant to ubonodin. Exponential-phase culture was plated at 10^8^ CFUs and 10 μL of 0-160 μM of ubonodin were spotted in the respective sectors. No zones of growth inhibition were observed after the spot-on-lawn LB plate was incubated at 30°C for ∼15 h.

## DISCUSSION

In this work, we have uncovered how the antibacterial lasso peptide ubonodin crosses the bacterial cell membranes as essential steps in its mechanism of action. Ubonodin is a narrow-spectrum antibacterial that inhibits transcription through inhibition of RNAP (Cheung-Lee et al., 2020). How does a compound with a highly conserved molecular target yet have confined bioactivity? Based on our observation that ubonodin is bioactive against select members of the Bcc, we developed a comparative genomics approach to predict the genes that differentiate susceptible from non-susceptible strains speculating that the capacity to internalize ubonodin determines susceptibility. This analysis revealed PupB, an OM transporter that putatively functions as a receptor for siderophore-iron complexes and that we have now shown can be hijacked by ubonodin for cellular entry (**Figure 8A**). By identifying PupB and demonstrating that it is necessary and sufficient to determine susceptibility to ubonodin, we have provided a molecular explanation for its spectrum of bioactivity. Our finding also provides an additional example of a lasso peptide that repurposes a siderophore receptor for OM transport, highlighting perhaps a universal strategy used by members of this RiPP class to breach the notoriously impenetrable bacterial OM.

**Figure 8.**
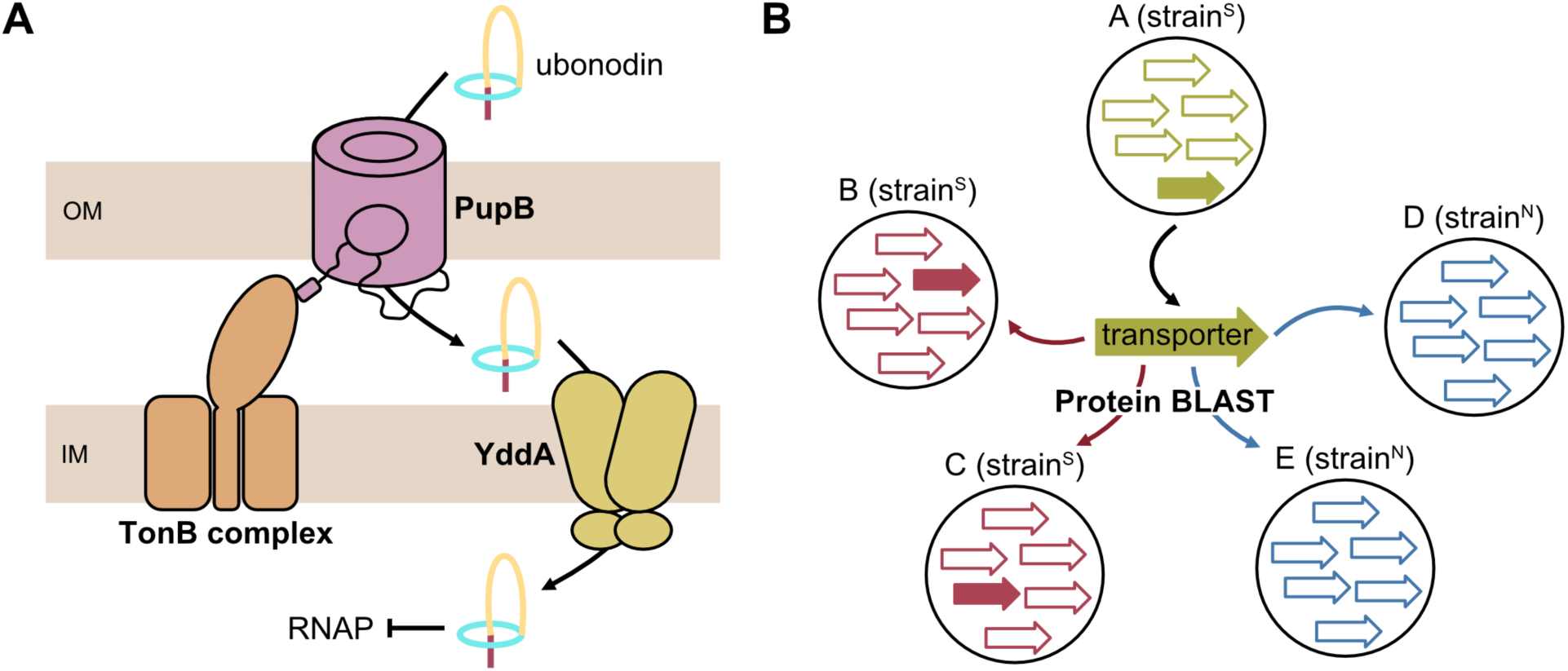
Model for the membrane transport pathway of ubonodin. (**A**) To cross the bacterial outer membrane (OM), ubonodin hijacks the siderophore receptor PupB native to certain Bcc strains. PupB functions in complex with its TonB activator. The inner membrane (IM) transporter of ubonodin is the putative ABC transporter YddA. (**B**) Comparative genomics was key to honing in on PupB and this approach should be broadly useful for investigating the mechanisms of action of other antimicrobial RiPPs with a defined susceptibility profile.

Lasso peptides that act as RNAP inhibitors comprise a major lasso peptide subclass and an emerging theme is that these peptides use TBDTs localized in the OM of target bacterial cells for initial cellular entry. Microcin J25 uses the *E. coli* TBDT FhuA for OM transport (Mathavan et al., 2014; Salomón and Farías, 1993). Klebsidin (Metelev et al., 2017) and microcin Y (Li et al., 2021) likewise use FhuA-like transporters. In these cases where the OM transporters were identified, the ability to readily isolate resistant mutants that have a mutation in the OM transporter or to predict a close homolog to *E. coli* FhuA was crucial to their identification. With ubonodin, we have observed a very low rate of bacterial resistance. While this property is desired for an antibiotic candidate, our difficulty in raising ubonodin-resistant *B. cepacia* mutants precluded straightforward identification of its OM transporter. Moreover, *B. cepacia* and multireplicon *Burkholderia* bacteria generally encode multiple homologs for a protein that is otherwise found only once in other bacteria (Mahenthiralingam et al., 2005). This unprecedented level of redundancy presented an additional challenge when we initially searched for TBDT homologs in *B. cepacia*, as there were 29 predicted TBDTs and the closest homolog was only ∼37% identical to *E. coli* FhuA. Without a clear candidate to assess, knocking out each TBDT one-by-one would be prohibitively time-consuming because genetic tools to manipulate *Burkholderia* are less developed (Barrett et al., 2008; Flannagan et al., 2008; Hogan et al., 2021, 2019; Shastri et al., 2017). Thus, we conceived an alternative strategy using comparative genomics to prioritize the *B. cepacia* TBDT candidates for further genetic studies to confirm their role in ubonodin transport. Ultimately, the putative *B. cepacia* TBDT PupB that we have established as the OM receptor for ubonodin is only ∼25% identical to *E. coli* FhuA. Our comparative genomics approach was key to honing in on PupB and we anticipate that it will be broadly useful in deciphering the mechanisms of actions of other lasso peptides and RiPPs (**Figure 8B**). We do note that a limitation of this strategy is that it cannot resolve whether strains encoding a susceptibility-defining gene actually express those genes, which might explain why *B. multivorans* AU15814 encodes *pupB* but is ubo^N^. For such exceptions, comparative transcriptomics on a panel of differentially-susceptible strains might be more appropriate.

Another key insight that linked siderophore receptors and specifically PupB to ubonodin transport was our finding that excess iron inhibits ubonodin bioactivity. RNA-seq profiling of *B. cepacia* indicated that excess iron represses *pupB* expression; fewer transporters would mean reduced cellular uptake of ubonodin and therefore bioactivity. Initial bioinformatic analysis suggested a signaling pathway involving the iron-sensing transcriptional repressor Fur. We had hoped to test that Fur directly controls *pupB* expression by deleting the *fur* gene in *B. cepacia* and showing that the mutant is hypersensitive to ubonodin, but we found that *fur* is essential. Although iron regulation in *Burkholderia* has been an area of significant interest, the specific role of *Burkholderia* Fur is poorly understood. A *fur* deletion mutant of *B. multivorans* has been reported, but this mutant has severe growth defects (Sato et al., 2017; Yuhara et al., 2008). Pleiotropic effects due to *fur* depletion were also observed in *P. aeruginosa* (Pasqua et al., 2017). While we could not delete *fur*, this line of investigation led us to another interesting observation. The operon that is predicted to contain *pupB* and to be under Fur regulation also encodes FecI and FecR, proteins that resemble an extracytoplasmic σ factor and an IM σ regulator pair. As we had discussed above, FecIR-type protein pairs cooperate with a special class of TBDTs that are structurally distinct from conventional TBDTs (Braun et al., 2003; Braun and Mahren, 2005; Visca et al., 2002). As PupB harbors the N-terminal extension domain found uniquely in this TBDT subclass, an intriguing question for future studies is whether PupB and FecIR form an iron-regulated signaling cascade in the native cellular context.

In addition to transcriptional regulation, as a TBDT PupB is also regulated through direct interaction with its TonB partner. Although *B. cepacia* encodes 4 TonB homologs, only TonB1 is required for ubonodin activity. Either the PupB-TonB1 interaction is highly specific or the other TonBs are not expressed under the experimental conditions, meaning that the Δ*tonB2*, Δ*tonB3*, and Δ*tonB4* mutants would effectively mimic WT. Given that expressing PupB alone in *E. coli* is sufficient for ubonodin susceptibility, we suspect there is some degree of promiscuity in the interaction of PupB with a TonB regulator. *E. coli* has one TonB regulator and it is only ∼28% identical to *B. cepacia* TonB1. Unless PupB is independently active in *E. coli*, it likely cooperates with *E. coli* TonB; expressing PupB in an *E. coli* Δ*tonB* background may provide an answer. By contrast, the interaction of PupB with ubonodin appears quite specific as no other *B. cepacia* or *E. coli* TBDTs can apparently substitute for its ubonodin transport function. Mapping the PupB-ubonodin binding interface will expose how the transporter engages its substrate, a subject for future studies.

Finally, while we show that OM transport is predictive of ubonodin susceptibility, bioactivity also depends on ubonodin’s ability to cross the IM as its intended cellular target is cytoplasmic. Based on its resemblance to *E. coli* SbmA, a known IM transporter for other RNAP-inhibiting lasso peptides (Cheung-Lee et al., 2019; Salomón and Farías, 1995), we honed in on the putative ABC transporter YddA as the ubonodin IM transporter in *B. cepacia*. The native biological function of YddA is not known but YddA is highly conserved in the Bcc strains examined herein. As both ubo^N^ and ubo^S^ Bcc strains encode near-identical YddA homologs, they all appear to be inherently capable of ubonodin IM transport. IM translocation is necessary but clearly not sufficient for ubonodin bioactivity, as further evidenced by the requirement for PupB in addition to a YddA-like transporter to observe activity against *E. coli*. Similar to our reconstitution of PupB function in *E. coli*, other groups have shown that expressing the native microcin J25 (Vincent et al., 2004) and klebsidin (Metelev et al., 2017) OM receptors in resistant strains without providing their native IM transporters is sufficient to achieve bioactivity. Thus, IM transporters capable of recognizing ubonodin and lasso peptides in general may be highly prevalent. Although SbmA homologs have been repeatedly associated with lasso peptide transport, how they interact with their cargo is poorly understood. In the case of *E. coli* SbmA-mediated transport of microcin J25, it is thought that SbmA makes specific contacts with the peptide and is powered by a proton gradient (Ghilarov et al., 2021). Unlike SbmA, which possesses the transmembrane but not nucleotide-binding domain of ABC transporters (Ghilarov et al., 2021; Runti et al., 2013), *B. cepacia* YddA is predicted to have a C-terminal ATP-binding domain. As such, YddA may use a transport mechanism distinct from SbmA, another intriguing question for future studies.

To date, hundreds of RiPPs have been discovered but for most the mechanism of action is still unknown, limiting their potential as clinical drugs. Before the genomic era, new RiPPs were typically found through bioactivity-guided studies, starting from isolation of a compound with antimicrobial activity to elucidation of its structure and target. Genomics-guided discovery of RiPPs has now overtaken activity-focused approaches, but our understanding of the function of these new compounds has not kept pace in part due to the lack of tools. The pipeline we have developed – from phenotype/drug susceptibility to computational genomics to target discovery – is generalizable to other RiPPs for which mechanism of action data are sorely needed.

## ACKNOWLEDGEMENTS

We thank Wei Wang and his staff at the Princeton Genomics Core Facility for advice on RNA isolation and for construction and sequencing of the RNA-seq libraries. We also thank Mark S. Thomas (University of Sheffield) for sharing his pSHAFT-series, p34E-Km, and p34E-TpTer plasmids and both him and Silvia T. Cardona (University of Manitoba) for useful advice in making *Burkholderia* mutants. We also thank Mohammad R. Seyedsayamdost for sharing the DH5α/pRK2013 helper strain for triparental mating and Zemer Gitai for the S17-1(λpir) and CC118(λpir) strains. Finally, we thank John LiPuma for sharing the Bcc clinical strains to test ubonodin susceptibility. Funding for this work was provided by the National Institutes of Health grant GM107036 to A.J.L.

## DATA AVAILABILITY STATEMENT

RNA-seq data (accession number PRJNA813900) can be found in the NCBI BioProject database.

## COMPETING INTERESTS

A.J.L. has applied for a patent covering the antimicrobial activity of ubonodin.

## MATERIALS AND METHODS

### Culture conditions

Unless otherwise indicated, all bacterial strains were cultured in Luria-Bertani Miller broth (10 g/L tryptone, 10 g/L NaCl, 5 g/L yeast extract). For spot-on-lawn assays, LB and M63 media (2 g/L (NH_4_)_2_SO_4_, 13.6 g KH_2_PO_4_, 0.2% D-glucose, 1 mM MgSO_4_, 0.04 g/L each of the 20 standard amino acids, 0.00005% thiamine) were used. For MIC liquid inhibition assays, cation-adjusted Mueller-Hinton II broth was also used (3 g/L beef extract, 17.5 g/L acid hydrolysate of casein, 1.5 g/L starch, 10 mg/L Mg^2+^, 20 mg/L Ca^2+^). *E. coli* strains were grown at 37°C overnight. *B. cepacia* strain ATCC 25416 was typically grown at 30°C for 1-2 days until distinct colonies appeared. All Bcc clinical isolates were grown at 32-35°C for 1-3 days until colonies appeared. Antibiotics were used at the following concentration for selection in *E. coli*: 100 μg/mL ampicillin, 25 μg/mL chloramphenicol, 50 μg/mL trimethoprim, 50 μg/mL kanamycin, and 50 μg/mL gentamicin. For selection in *B. cepacia*, antibiotics were used at the following concentrations: 300 μg/mL kanamycin, 50-100 μg/mL trimethoprim, and 50 μg/mL chloramphenicol.

### Strain construction

The list of strains used in this study can be found in **Supplementary file 4**. Chemical transformation was used to introduce plasmids into *E. coli* XL1-Blue Mix & Go competent cells (Zymo Research, cat. no. T3002) or *E. coli* DH5α chemically competent cells. Electroporation was used to introduce pir-dependent plasmids into electrocompetent *E. coli* CC118(λpir) and *E. coli* S17-1(λpir) cells. Electroporation was also used to introduce plasmids into electrocompetent *E. coli* BL21 and BL21(DE3) Δ*slyD* cells. Abbreviations used in this and the plasmid construction sections: Cm^R^, chloramphenicol resistance; Gent^R^, gentamicin resistance, Tp/TpTer/Tp^R^, trimethoprim resistance; Km/Km^R^, kanamycin resistance.

#### TD042

[*B. cepacia* ATCC 25416 ΔGGFLHMPP_06959 (*tonB3*)::TpTer]. The donor strain *E. coli* S17-1(λpir) propagating the pTD005-[pSHAFT3-Δ*tonB3*::TpTer] suicide plasmid was biparentally mated into recipient strain *B. cepacia* ATCC 25416. Tp^R^Gent^R^ transconjugant colonies were screened to identify double-crossover mutants which are also Cm^S^ and further confirmed by colony PCR using primer pairs oTD045-oTD046 and oTD045-oTD043.

#### TD075

[*B. cepacia* ATCC 25416 ΔGGFLHMPP_00610 (*fur*)::Km; pSCrhaB2-*fur*]. This strain was made in two steps by first introducing the *fur* expression plasmid and then replacing chromosomal *fur* with a Km^R^ marker. The donor strain *E. coli* DH5α propagating pTD013-[pSCrhaB2-*fur*] and helper strain *E. coli* DH5α/pRK2013 were triparentally mated with recipient strain *B. cepacia* ATCC 25416 to transfer the expression plasmid. Tp^R^Gent^R^ transconjugant colonies were checked by colony PCR using primer pairs oTD035-oTD036 to confirm transfer of pTD013. Next, the donor strain *E. coli* S17-1(λpir) propagating the pTD016-[pTD014-Δ*fur*::Km] suicide plasmid was biparentally mated into recipient strain *B. cepacia* ATCC 25416; pSCrhaB2-*fur*. Km^R^Gent^R^ transconjugant colonies were screened to identify double-crossover mutants which are also Cm^S^ and further confirmed by colony PCR using primer pairs oTD062-oTD063 and oTD063-oTD091. Strain TD075 needed to be propagated on LB selection plates supplemented with 0.02-0.2% L-rhamnose as *fur* is essential.

#### TD078

[*B. cepacia* ATCC 25416 ΔGGFLHMPP_07362 (*tonB4*)::Km]. The donor strain *E. coli* S17-1(λpir) propagating the pTD017-[pSHAFT3-Δ*tonB4*)::Km] suicide plasmid was biparentally mated into recipient strain *B. cepacia* ATCC 25416. Km^R^Gent^R^ transconjugant colonies were screened to identify double-crossover mutants which are also Cm^S^ and further confirmed by colony PCR using primer pairs oTD089-oTD090 and oTD090-oTD091.

#### TD080

[*B. cepacia* ATCC 25416; pSCrhaB2]. The donor strain *E. coli* DH5α propagating pSCrhaB2 and helper strain *E. coli* DH5α/pRK2013 were triparentally mated with recipient strain *B. cepacia* ATCC 25416 to transfer the empty vector. Tp^R^Gent^R^ transconjugant colonies were checked by colony PCR using primer pairs oTD035-oTD036 to confirm transfer of pSCrhaB2.

#### TD090

[*B. cepacia* ATCC 25416 ΔGGFLHMPP_04892 (*tonB2*)::TpTer]. The donor strain *E. coli* S17-1(λpir) propagating the pTD022-[pTD014-Δ*tonB2*::TpTer] suicide plasmid was biparentally mated into recipient strain *B. cepacia* ATCC 25416. Tp^R^Gent^R^ transconjugant colonies were screened to identify double-crossover mutants which are also Cm^S^ and further confirmed by colony PCR using primer pairs oTD132-oTD133 and oTD132-oTD043.

#### TD091

[*B. cepacia* ATCC 25416 ΔGGFLHMPP_02544 (*tonB1*)::Km; pSCrhaB2-*tonB1*]. This strain was made in two steps by first introducing the *tonB1* expression plasmid and then replacing chromosomal *tonB1* with a Km^R^ marker. The donor strain *E. coli* DH5α propagating pTD020-[pSCrhaB2-*tonB1*] and helper strain *E. coli* DH5α/pRK2013 were triparentally mated with recipient strain *B. cepacia* ATCC 25416 to transfer the expression plasmid. Tp^R^Gent^R^ transconjugant colonies were checked by colony PCR using primer pairs oTD035-oTD036 to confirm transfer of pTD020. Next, the donor strain *E. coli* S17-1(λpir) propagating the pTD019-[pTD014-Δ*tonB1*::Km] suicide plasmid was biparentally mated into recipient strain *B. cepacia* ATCC 25416; pSCrhaB2-*tonB1*. Km^R^Gent^R^ transconjugant colonies were screened to identify double-crossover mutants which are also Cm^S^ and further confirmed by colony PCR using primer pairs oTD118-oTD119 and oTD118-oTD091. Strain TD091 was propagated on LB selection plates supplemented with 0.02% L-rhamnose as *tonB1* is important for healthy cell growth.

#### TD103

[*B. cepacia* ATCC 25416 ΔGGFLHMPP_01381 (*pupB*)::Km]. The donor strain *E. coli* S17-1(λpir) propagating the pTD023-[pTD014-Δ*pupB*::Km] suicide plasmid was biparentally mated into recipient strain *B. cepacia* ATCC 25416. Km^R^Gent^R^ transconjugant colonies were screened to identify double-crossover mutants which are also Cm^S^ and further confirmed by colony PCR using primer pairs oTD151-oTD152.

#### TD118

[*B. cepacia* ATCC 25416 ΔGGFLHMPP_01381 (*pupB*)::Km; pSCrhaB2-APZ15_10615 (*pupB*)]. This strain was made in two steps by first introducing the *pupB* expression plasmid and then replacing chromosomal *pupB* with a Km^R^ marker. The donor strain *E. coli* DH5α propagating pTD029-[pSCrhaB2-*pupB*] and helper strain *E. coli* DH5α/pRK2013 were triparentally mated with recipient strain *B. cepacia* ATCC 25416 to transfer the expression plasmid. Tp^R^Gent^R^ transconjugant colonies were checked by colony PCR using primer pairs oTD035-oTD036 to confirm transfer of pTD029. Next, the donor strain *E. coli* S17-1(λpir) propagating the pTD023-[pTD014-Δ*pupB*::Km] suicide plasmid was biparentally mated into recipient strain *B. cepacia* ATCC 25416; pSCrhaB2-*pupB*. Km^R^Gent^R^ transconjugant colonies were screened to identify double-crossover mutants which are also Cm^S^ and further confirmed by colony PCR using primer pairs oTD151-oTD152.

#### TD12

[*B. cepacia* ATCC 25416 ΔGGFLHMPP_00373 (*yddA*)::TpTer]. The donor strain *E. coli* S17-1(λpir) propagating the pTD032-[pTD014-Δ*yddA*::TpTer] suicide plasmid was biparentally mated into recipient strain *B. cepacia* ATCC 25416. Tp^R^Gent^R^ transconjugant colonies were screened to identify double-crossover mutants which are also Cm^S^ and further confirmed by colony PCR using primer pairs oTD180-oTD179 and oTD180-oTD043.

### Plasmid construction

The list of plasmids and oligonucleotides used in this study can be found in **Supplementary file 5** and **Supplementary file 6**, respectively. Plasmids were cloned using *E. coli* XL1-Blue and *E. coli* DH5α for general purposes, but pir-dependent pSHAFT3-based plasmids were cloned using *E. coli* CC118(λpir). The pWC99 plasmid for ubonodin purification was overexpressed in *E. coli* BL21. Plasmids for *B. cepacia* PupB and/or TonB1 overexpression were freshly transformed into *E. coli* BL21(DE3) Δ*slyD*. When needed, bacterial genomic DNA was extracted using the DNeasy Blood & Tissue Kit (QIAGEN, cat. no. 69504) according to manufacturer’s protocol. The Q5® High-Fidelity DNA Polymerase and buffers (NEB, cat. no. M0491L) were used for PCR. Other standard cloning reagents include T4 DNA ligase (NEB), restriction enzymes (NEB), Zymoclean Gel DNA Recovery Kits (Zymo Research), and QIAprep Spin Miniprep Kits (QIAGEN). All plasmids were sequenced using the GENEWIZ/Azenta Life Sciences sequencing service to confirm the correct insert was cloned.

#### pTD00

[pSHAFT3-ΔGGFLHMPP_06959 (*tonB3*)::TpTer]. The 692-bp sequence upstream of the *tonB3* ATG start codon was PCR-amplified using primer pairs oTD026-oTD027 and ATCC 25416 gDNA template. The PcS-TpTer cassette was PCR-amplified using primer pairs oTD029-oTD030 and p34E-TpTer plasmid template. The 783-bp sequence downstream of *tonB3* Ser227 codon was PCR-amplified using primer pairs oTD028-oTD025 and ATCC 25416 gDNA template. The 3 PCR products were stitched together with T4 DNA ligase and cloned between the EcoRI and XbaI sites of pSHAFT3.

#### pTD013

[pSCrhaB2-GGFLHMPP_00610 (*fur*)]. GGFLHMPP_00610 [Met1-His142] was PCR-amplified using primer pairs oTD083-oTD084 and ATCC 25416 gDNA template. The PCR product was cloned between the NdeI and BamHI sites of pSCrhaB2.

#### pTD014

[pSHAFT3-Amp^R^ Cyt717Ade-BsaI-PGphD-mCherry-BsaI]. This plasmid was made in two steps. First, the Amp^R^ [Ile140-Gly242] sequence was PCR-amplified using primer pairs oTD064-oTD065 and pSHAFT3 plasmid template. Primer oTD064 mutates out the native BsaI within the pSHAFT3 Amp^R^ sequence. The PCR product was cloned between the BsaI and PvuI sites of pSHAFT3. Next, the PGphD promoter was PCR-amplified using primer pairs oTD076-oTD078 and pYTK001 plasmid template, a plasmid made available on Addgene (#65108) from John Dueber (Lee et al., 2015). The mCherry sequence was PCR-amplified using primer pairs oTD079-oTD080 and pET28a-mCherry-CNA35 plasmid template, a plasmid made available on Addgene (#61607) from Maarten Merkx (Aper et al., 2014). Overlap PCR was performed to stitch together the PGphD and mCherry products, and the resultant product was cloned between the EcoRI and XbaI sites of the plasmid generated in the first step.

#### pTD016

[pTD014-ΔGGFLHMPP_00610 (*fur*)::Km]. The 702-bp sequence upstream of the *fur* Thr5 codon was PCR-amplified using primer pairs oTD094-oTD095 and ATCC 25416 gDNA template. The PcS-Km cassette was PCR-amplified using primer pairs oTD098-oTD099 and p34E-Km plasmid template. The 671-bp sequence downstream of the *fur* Arg140 codon was PCR-amplified using primer pairs oTD096-oTD097 and ATCC 25416 gDNA template. The 3 PCR products were cloned between the EcoRI and XbaI sites of pTD014 via Golden Gate cloning.

#### pTD017

[pSHAFT3-ΔGGFLHMPP_07362 (*tonB4*)::Km]. The 742-bp sequence upstream of the *tonB4* Met1 start codon was PCR-amplified using primer pairs oTD071-oTD104 and ATCC 25416 gDNA template. The PcS-Km cassette was PCR-amplified using primer pairs oTD106-oTD107 and p34E-Km plasmid template. The 716-bp sequence downstream of the *tonB4* Phe223 codon was PCR-amplified using primer pairs oTD105-oTD068 and ATCC 25416 gDNA template. The 3 PCR products were stitched together using overlap PCR and cloned between the EcoRI and XbaI sites of pSHAFT3.

#### pTD019

[pTD014-ΔGGFLHMPP_02544 (*tonB1*)::Km]. The 615-bp sequence upstream of the *tonB1* Pro3 codon was PCR-amplified using primer pairs oTD114-oTD115 and ATCC 25416 gDNA template. The PcS-Km cassette was PCR-amplified using primer pairs oTD098-oTD099 and p34E-Km plasmid template. The 588-bp sequence downstream of the *tonB1* stop codon was PCR-amplified using primer pairs oTD116-oTD117 and ATCC 25416 gDNA template. The 3 PCR products were cloned between the EcoRI and XbaI sites of pTD014 via Golden Gate cloning.

#### pTD020

[pSCrhaB2-GGFLHMPP_02544 (*tonB1*)]. GGFLHMPP_02544 [Met1-Asp226] was PCR-amplified using primer pairs oTD120-oTD121 and ATCC 25416 gDNA template. The PCR product was cloned between the NdeI and BamHI sites of pSCrhaB2.

#### pTD022

[pTD014-ΔGGFLHMPP_04892 (*tonB2*)::TpTer]. The 670-bp sequence upstream of the *tonB2* Met1 codon was PCR-amplified using primer pairs oTD128-oTD129 and ATCC 25416 gDNA template. The PcS-TpTer cassette was PCR-amplified using primer pairs oTD029-oTD030 and p34E-TpTer plasmid template. The 625-bp sequence downstream of the *tonB2* Asp269 codon was PCR-amplified using primer pairs oTD130-oTD131 and ATCC 25416 gDNA template. The 3 PCR products were cloned between the EcoRI and XbaI sites of pTD014 via Golden Gate cloning.

#### pTD023

[pTD014-ΔGGFLHMPP_01381 (*pupB*)::Km]. The 686-bp sequence upstream of the *pupB* Val2 codon was PCR-amplified using primer pairs oTD134-oTD135 and ATCC 25416 gDNA template. The PcS-Km cassette was PCR-amplified using primer pairs oTD098-oTD099 and p34E-Km plasmid template. The 724-bp sequence downstream of the *pupB* Leu840 codon was PCR-amplified using primer pairs oTD136-oTD137 and ATCC 25416 gDNA template. The 3 PCR products were cloned between the EcoRI and XbaI sites of pTD014 via Golden Gate cloning.

#### pTD027

[pRSFDuet-1-GGFLHMPP_02544 (*tonB1*)]. GGFLHMPP_02544 [Met1-Asp226] was PCR-amplified using primer pairs oTD168-oTD169 and ATCC 25416 gDNA template. The PCR product was cloned between the NcoI and HindIII sites of pRSFDuet-1.

#### pTD029

[pSCrhaB2-APZ15_10615 (*pupB*)]. APZ15_10615 [Met1-Phe858] was PCR-amplified using primer pairs oTD173-oTD174 and ATCC 25416 gDNA template. The PCR product was cloned between the NdeI and HindIII sites of pSCrhaB2.

#### pTD030

[pRSFDuet-1-GGFLHMPP_02544 (*tonB1*)-APZ15_10615 (*pupB*)]. APZ15_10615 [Met1-Phe858] was PCR-amplified using primer pairs oTD173-oTD170 and ATCC 25416 gDNA template. The PCR product was cloned between the NdeI and AvrII sites of pTD027.

#### pTD031

[pRSFDuet-1-APZ15_10615 (*pupB*)]. APZ15_10615 [Met1-Phe858] was PCR-amplified using primer pairs oTD173-oTD170 and ATCC 25416 gDNA template. The PCR product was cloned between the NdeI and AvrII sites of pRSFDuet-1.

#### pTD032

[pTD014-ΔGGFLHMPP_00373 (*yddA*)::TpTer]. The 634-bp sequence upstream of the *yddA* Met1 codon was PCR-amplified using primer pairs oTD175-oTD176 and ATCC 25416 gDNA template. The PcS-TpTer cassette was PCR-amplified using primer pairs oTD029-oTD030 and p34E-TpTer plasmid template. The 605-bp sequence downstream of the *yddA* Val581 codon was PCR-amplified using primer pairs oTD177-oTD178 and ATCC 25416 gDNA template. The 3 PCR products were cloned between the EcoRI and XbaI sites of pTD014 via Golden Gate cloning.

### Biparental mating

Biparental mating was used to transfer pSHAFT3-based suicide plasmids (Shastri et al., 2017) from *E. coli* into *B. cepacia* for allelic replacement. Briefly, 50 μL each of overnight cultures of the donor strain propagating the suicide plasmid and the recipient strain were mixed. The cell mixture was centrifuged at 6000 xg for 2 min at room temperature, and the cell pellet was resuspended with 50 μL of LB and spotted onto an LB agar plate without antibiotic selection. The plate was incubated face up at 30°C overnight. The next day, the mating spot was scraped up with an inoculation loop and resuspended in 200 μL of sterile 1X PBS buffer. The resuspended cell mixture was plated on the appropriate selection plates to select for transconjugants. Gentamicin was used to select against the donor strain. Transconjugant colonies were picked and patched to screen for Cm^S^ indicating double-crossover mutants that have lost the wild-type allele.

### Triparental mating

Triparental mating was used to transfer pSCrhaB2-based plasmids from *E. coli* into *B. cepacia*. Our study adapted a previously published protocol for triparental mating (Aubert et al., 2014). Briefly, 50 μL each of overnight cultures of the donor strain propagating the cargo plasmid, the helper strain *E. coli* DH5α/pRK2013 (Figurski and Helinski, 1979), and the recipient strain were mixed. The cell mixture was centrifuged at 6000 xg for 2 min at room temperature, and the cell pellet was resuspended with 50 μL of LB and spotted onto an LB agar plate without antibiotic selection. The plate was incubated face up at 30°C overnight. The next day, the mating spot was scraped up with an inoculation loop and resuspended in 200 μL of sterile 1X PBS buffer. The resuspended cell mixture was plated on the appropriate selection plates to select for transconjugants. Gentamicin was used to select against the donor strain.

### Expression and purification of ubonodin

Ubonodin was overexpressed and purified as previously reported (Cheung-Lee et al., 2020). Briefly, BL21 propagating pWC99 was grown in M9 media (6.4 mg/L Na_2_HPO_4_ · 7 H_2_O, 1.5 mg/L KH_2_PO_4_, 0.5 mg/L NH_4_Cl, 0.25 mg/L NaCl, 0.15 μg/mL CaCl_2_, 1 mM MgSO_4_, 0.2% D-glucose, 0.04 g/L each of the 20 standard amino acids, 0.00005% thiamine) and ubonodin expression was induced by the addition of 1 mM IPTG. Cultures were grown for ∼20 h at 20°C following induction and then centrifuged to harvest the supernatant. The supernatant was extracted through a Strata C8 column, eluted with 100% methanol, and dried by rotovapping. The dried extract was resuspended with 25% acetonitrile/75% water and further purified by semi-preparative reverse-phase HPLC using an Agilent Technologies 1200 series instrument fitted with a Zorbax 300SB-C18 column (9.4 mm x 250 mm, 5 μm). Purified ubonodin was lyophilized and resuspended in water. The purity was checked by LC-MS and the concentration was measured using the A280 Nanodrop ND-1000 Spectrophotometer reading.

### Cladogram construction

To construct the cladogram of Bcc strains tested for ubonodin susceptibility, first the allelic profile for 7 Bcc housekeeping genes (*atpD, gltB, gyrB, recA, lepA, phaC*, and *trpB*) and sequence type of each strain were retrieved from the *Burkholderia* Genome Database (Winsor et al., 2008). Then, the allelic profile and sequence type were entered into the PubMLST website (Public databases for molecular typing and microbial genome diversity) to obtain the concatenated sequences of fragments of the 7 genes listed above. Finally, the concatenated sequences for all Bcc strains were aligned using Clustal Omega to generate the cladogram (Sievers et al., 2011).

### RNA polymerase protein sequence alignment

To generate a percent sequence identity matrix comparing the RNA polymerase β and β’ subunits encoded by the Bcc strains tested for ubonodin susceptibility, first the RpoB and RpoC sequences for each strain were retrieved from NCBI. The concatenated RpoBC sequences were then aligned using Clustal Omega (Sievers et al., 2011).

### Spot-on-lawn assay

A standard spot-on-lawn assay to assess bacterial susceptibility to ubonodin was carried out as previously reported (Cheung-Lee et al., 2020) with a few changes denoted here. Overnight cultures of test strains were diluted 1:100 into fresh LB with antibiotics as needed the following day and grown at the appropriate temperature with shaking to an OD_600_ of ∼0.4-0.6. Once the cultures reached mid-exponential phase, the volume equivalent to 10^8^ CFUs (assuming OD_600_ = 1 contains 10^9^ CFUs/mL) was added to 10 mL of melted LB or M63 soft agar (0.65% agar). The cell and agar mixture was poured onto 10 mL of LB or M63 base agar (1.5% agar, M63 base agar lacks amino acids) and the plate was left to dry in the biosafety cabinet with the lid slightly ajar. When needed, IPTG or L-rhamnose was also added to the soft agar to induce protein expression. M63 was used for the Bcc clinical isolates and *E. coli* BL21(DE3) Δ*slyD* overexpression strains, whereas the ATCC 25416 strains were assessed on LB media. The ubonodin MIC of *B. cepacia* WT is similar on M63 and LB. Two-fold serial dilutions of ubonodin (0-160 μM) were prepared using sterile ultrapure water and 10 μL of each dilution was spotted onto the dried lawn of cells in their respective sector. The spots were left to dry in the biosafety cabinet once again with the lid slightly ajar. The plates were incubated ∼15 h at the appropriate temperature and imaged the next day using the Bio Rad ChemiDoc XRS Gel Imaging System under epi white illumination and the Quantity One 4.6.6 imaging software. Images were processed using FIJI. Spot-on-lawn assays were performed with at least 2 biological replicates (independent cultures) for each tested ATCC 25416 strain and the Bcc clinical isolates AU0158 and AU15814.

### Spot dilution assay

To assess cell viability by spot dilution assay, overnight cultures of test strains were diluted 1:100 into fresh LB with antibiotics as needed the following morning and grown at 30°C with shaking to an OD_600_ of ∼0.4-0.6. Once the cultures reached mid-exponential phase, all cultures for the set of strains being tested were normalized to the OD_600_ of the least dense culture using LB. From this 10^0^ starting dilution, ten-fold serial dilutions were prepared using LB. Five μL of each dilution was spotted onto the test media. The spots were left to dry in the biosafety cabinet with the lid slightly ajar. The plates were incubated ∼15 h at the appropriate temperature and imaged the next day using the Bio Rad ChemiDoc XRS Gel Imaging System under epi white illumination and the Quantity One 4.6.6 imaging software. Images were processed using FIJI. Spot dilution assays were performed with at least 2 biological replicates (independent cultures) for each tested strain.

### Liquid inhibition assay and bacterial growth rate analysis

To assess the endpoint MIC of ubonodin by liquid inhibition assay, overnight cultures of test strains were diluted 1:100 into fresh LB with antibiotics as needed and grown at 30°C with shaking to an OD_600_ of ∼0.4-0.6. Once the cultures reached mid-exponential phase, they were diluted to an OD_600_ of 0.001 in the test media. Two-fold serial dilutions of ubonodin (0-32 μM) were prepared in test media and 50 μL of each dilution was dispensed into an untreated, sterile 96-well plate. The ubonodin dilutions were topped off with an equivalent volume (50 μL) of the 0.001 OD_600_ cells, bringing the final OD_600_ to 0.0005 in each well and the range of ubonodin concentrations to 0-16 μM. The 96-well plate was covered with a matching lid and incubated with shaking at 30°C for ∼16 h. The endpoint OD_600_ was measured using a BioTek Synergy 4 plate reader and blanked to a test media only control to calculate the MIC of ubonodin. When needed, the test medium was supplemented with L-rhamnose to induce protein expression. Both Mueller-Hinton Broth II and LB were used as test media for liquid inhibition assays; the media used in a certain assay is indicated within the corresponding figure caption.

To assess bacterial growth over time, exponential-phase LB cultures of test strains inoculated from a scoop of freshly-streaked colonies were diluted to a final OD_600_ of 0.001 in 150 μL of test media in a 96-well plate. The plate was covered with a matching lid and incubated with shaking at 30°C in a BioTek Synergy 4 plate reader. OD_600_ measurements were recorded at 20-min intervals for 20-24 h.

### Protein BLAST-based comparative genomics

Protein BLAST-based comparative genomics was performed using the standalone NCBI BLAST+ suite installed on a local machine (Camacho et al., 2009). For each strain, genome annotation files in GenBank format were downloaded from NCBI. The GenBank files were converted to FASTA format using the custom Python script gbk_to_fasta.py adapted from source code available through the University of Warwick at the following web link: https://warwick.ac.uk/fac/sci/moac/people/students/peter_cock/python/genbank2fasta/. Then, using the makeblastdb program within the BLAST+ suite, a protein BLAST database was built from each of the FASTA files.

The input sequences for the next step of analysis were retrieved by first predicting TBDT homologs in *B. cepacia* ATCC 25416. Using the *E. coli* FhuA sequence and 34 known and predicted *P. aeruginosa* TBDT sequences (Luscher et al., 2018) as queries, protein BLAST search was performed against the ATCC 25416 protein BLAST database to identify 29 *B. cepacia* TBDT homologs. A high e-value cutoff of 1000 was used in this search to maximize retrieval of any potential homologs. Using the BLAST+ suite blastp program, the 29 predicted TBDT homologs encoded by *B. cepacia* ATCC 25416 then served as queries for protein BLAST against each of the custom-built Bcc protein BLAST database generated in the first step. This process identifies the closest homolog to each predicted ATCC 25416 TBDT in each Bcc strain. Finally, to identify patterns of TBDT conservation across all strains, the custom blast_comparison_genes.py Python script was implemented. Briefly, this script calculates the percentage of positive-scoring matches (ppos; percent protein sequence similarity for the alignment of the query TBDT and subject TBDT hit) normalized by multiplying to the length of the alignment (qcovhsp; query coverage per high-scoring segment pair) for each query-top subject hit pair. A clustered 29 × 11 (TBDTs × BCC strains) heat map matrix was generated based on these normalized scores to visualize how conserved each predicted *B. cepacia* TBDT is across the other Bcc strains. This BLAST matrix analysis was adapted from a large-scale genomics study on *Burkholderia* strains (Ussery et al., 2009). GraphPad Prism version 9.3.1 was used to make all plots. All custom Python scripts used for this study were uploaded to the Link lab Github page at the following web link: https://github.com/ajlinklab/PupB.

### Ubonodin cellular uptake assay

The endpoint concentration of ubonodin in the cell pellet was analyzed as a measure of cellular uptake. Overnight cultures of test strains were diluted 1:100 into fresh LB and grown at 30°C with shaking to an OD_600_ of ∼0.4-0.6. The cultures were diluted to an OD_600_ of 0.01 in M63 broth and 3 × 150 μL aliquots were dispensed into separate wells of an untreated, sterile 96-well plate. One well was left untreated, the second well contained 1 μM ubonodin, and the third well contained 2 μM ubonodin. The plate was covered with a matching lid and incubated with shaking at 30°C for ∼16 h. The endpoint OD_600_ was measured using a BioTek Synergy 4 plate reader and blanked to an M63 media only control. The content of each well was transferred to a 1.5-mL microcentrifuge tube and the cell pellet was harvested at 6000 xg for 2 min at room temperature. The supernatant was removed and the cell pellet was washed once with 200 μL of sterile 1X PBS before a second spin to remove the wash fraction.

We adapted a previously published bacterial cell lysis protocol to release intracellular metabolites for LC-MS analysis (Pinu et al., 2017). The cell pellet was resuspended in 200 μL of cold 100% methanol. The cell resuspension was transferred to dry ice for 5 min, followed by thawing at room temperature and vortexing for 30 sec. This freeze-thaw-vortex step was repeated 3 times in total and the lysate was spun down at 10000 xg for 5 min at room temperature to harvest the supernatant and pellet out the cell debris. A second extraction step was performed to maximize cell lysis. The cell debris pellet was resuspended with another 200 μL of cold 100% methanol and subjected to another round of freeze-thaw-vortex before centrifugation. The supernatant from this second extraction was combined with the first extraction. To evaporate off the methanol of all samples in parallel, the methanol extracts were left in the fume hood at room temperature for ∼24 h with the cap open.

The dried extracts were resuspended with 75 μL of sterile ultrapure water and centrifuged at 10000 xg for 15 min at room temperature to pellet out debris. Thirty-five μL of the supernatant was transferred to an LC-MS vial fitted with a sample insert and 25 μL was injected onto the LC-MS for analysis. LC-MS analysis was performed using a Zorbax 300SB-C18 column (2.1 mm × 50 mm, 3.5 μm) installed on an Agilent 1260 Infinity II system in line with an Agilent 6530 Q-TOF mass spectrometer. The LC-MS gradient used for separation was: 0.5 mL/min 90% solvent A (H_2_O, 0.1% formic acid) and 10% solvent B (acetonitrile, 0.1% formic acid) for 1 min followed by a linear gradient to 50% solvent B over 19 min, then a linear gradient to 90% solvent B over 5 min, and a linear gradient back to 10% solvent B over 5 min. As ubonodin undergoes oxidation (Cheung-Lee et al., 2020) especially upon air drying in the fume hood, multiply-oxidized ubonodin species were observed by LC-MS. To calculate the intracellular concentration of ubonodin for each sample, the sum of the extracted ion counts of non-oxidized ubonodin and the ubonodin +16/32/48/64 species was calculated for the total sample volume of 75 μL and normalized to the final OD_600_.

### Extraction, preparation, and sequencing of total RNA

*B. cepacia* ATCC 25416 was freshly streaked out on LB agar. For each condition, three biological replicates derived from separate colonies were prepared. Single colonies were inoculated into LB ± 1 mM FeCl_3_ (ferric chloride stock was made fresh in water and filter-sterilized) and grown overnight at 30°C with shaking. The next morning, the overnight cultures were diluted 1:100 into fresh LB ± 1 mM FeCl_3_ and grown at 30°C with shaking to an OD_600_ of ∼0.5-0.6. The volume of mid-exponential phase culture to achieve 5 × 10^8^ CFUs was calculated (assuming OD_600_ of 1.0 yields 10^9^ CFUs/mL) and added to 2 volumes of RNAprotect Bacteria Reagent (QIAGEN, cat. no. 74124). The mixture was immediately vortexed for 5 sec, incubated at room temperature for 5 min, and then centrifuged at 5000 xg for 10 min at room temperature. The supernatant was decanted and the cell pellet was stored at -80°C until RNA extraction.

Total RNA was extracted from RNAprotect-stabilized cell pellets using the RNeasy Protect Kit (QIAGEN, cat. no. 74124) according to manufacturer’s protocol. Briefly, the frozen cell pellets were thawed at room temperature and cell lysis was achieved using enzymatic lysis with 200 μL of 1X TE lysis buffer (30 mM Tris-HCl, 1 mM EDTA, pH 8.0) containing 1 mg/mL of lysozyme and supplemented with 10 μL of Proteinase K (QIAGEN, cat. no. 19131). The mixture was incubated on a shaker at room temperature for 30 min with intermittent vortexing, 700 μL of Buffer RLT was added, the mixture was vortexed, and 500 μL of 100% ethanol was added and mixed. Total RNA was then purified from the bacterial cell lysate using the RNeasy Mini Kit protocol and eluted with 2 × 35-40 μL of nuclease-free water. The miniprepped RNA was further treated with in-solution rigorous TURBO DNase treatment using the TURBO DNA-free™ Kit (Thermo Fisher Scientific, cat. no. AM1907) to remove contaminant genomic DNA according to manufacturer’s protocol. The concentration and integrity of purified RNA were estimated using a NanoDrop ND-1000 Spectrophotometer and a 2% ethidium bromide non-denaturing agarose gel observing for intact rRNA bands, respectively. RNA samples were submitted to the Princeton Genomics Core Facility for library construction, quality control check, and sequencing on an Illumina NovaSeq 6000 Illumina sequencing platform. The samples were depleted of ribosomal RNA and checked on a Bioanalyzer prior to sequencing.

Analysis of RNA-Seq results was carried out in collaboration with research-computing staff at the Princeton University Lewis-Sigler Institute for Integrative Genomics using Galaxy. The *B. cepacia* ATCC 25416 genome assembly (ASM141149v1; GenBank assembly accession: GCA_001411495.1) was downloaded from NCBI in fna and gtf format; the gtf file was edited to avoid tool issues on Galaxy. Forward and reverse sequences were uploaded to and demultiplexed on the Princeton HTSeq database system and transferred to the Princeton Galaxy instance. Read quality was assessed using FastQC (Galaxy Version 0.72), Read Distribution (Galaxy Version 2.6.4.1), BAM/SAM Mapping Stats (Galaxy Version 2.6.4), IdxStats (Galaxy Version 2.0.2), and Gene Body Coverage (Galaxy Version 2.6.4.3). Top over-represented sequences and ribosomal RNA content was assessed using in-house Galaxy workflows. Quality control stats were viewed using a MultiQC (Galaxy Version 1.8+galaxy0) report. Sequences were aligned using Burrows-Wheeler Alignment (BWA; Galaxy Version 0.7.17.4) (Li and Durbin, 2009). Reads aligning to genes according to NCBI’s gene annotations were counted using featureCounts (Galaxy Version 1.6.4+galaxy1). Iron-treated and untreated samples were then compared using DESeq2 (Galaxy Version 2.11.40.6+galaxy1) (Love et al., 2014), which generated QC plots, rLog normalized counts, and differential expression data, including adjusted p-values to account for multiple testing with the Benjamini-Hochberg procedure which controls false discovery rate (FDR).

### Computational prediction of Fur binding sites

To predict Fur DNA-binding sites in the *B. cepacia* ATCC 25416 genome, the custom Python script PWMmodel.py was implemented on the Princeton Della server. This analysis was adapted from bioinformatics methods used for identification of regulatory elements (Wasserman and Sandelin, 2004). It first required a set of known Fur DNA-binding site sequences, which are in general well-conserved. Seven experimentally-determined 19-bp Fur binding sequences from *P. aeruginosa* were used as the known set (Dudek and Jahn, 2021; Hassett et al., 1997; Ochsner et al., 2000, 1995). Alignment of these 7 sequences using the online MEME tool generated a sequence logo summarizing the Fur consensus sequence (Bailey et al., 2009). From these 7 sequences, a position frequency matrix (PFM) was also generated by calculating the number of times each nucleotide (A, C, G, T) was found at each position within the 19-bp sequence. The PFM captures the nucleotide characteristics at each position. The PFM was then converted to a position weight matrix (PWM) by calculating the probability of observing a particular nucleotide at a specific position normalized to the expected background probability. The *B. cepacia* ATCC 25416 genome is ∼67% GC-rich, so 16.5% was expected for A/T each and 33.5% was expected for G/C each as the background probabilities. Using the PWM, any 19-bp sequence can be scored for how similar it is to the expected Fur binding site sequence. This is done by summing the individual PWM scores corresponding to an observed nucleotide at each consecutive position. The PWM built from the 7 known Fur binding sites was used to scan the ATCC 25416 genome for high-scoring Fur boxes. The scan made use of a 19-bp sliding window, which shifted by 1 nucleotide in each iteration, on both the forward and reverse strands. As many genome fragments are analyzed and most produce low match scores, this study filtered for Fur box hits that are found proximal to, but not within, and facing the same direction as a coding DNA sequence (CDS). It also filtered for Fur box hits in terms of information content, a measure of conservation at each position. To generate histograms of the match scores for all or a subset of scanned fragments, the custom Python script PWM-annotator.py was implemented on the Della server. All custom scripts used in this analysis can be found on the Link lab Github page.

## Notes

### Competing Interest Statement

The authors have declared no competing interest.

## REFERENCES

Andrews SC, Robinson AK, Rodríguez-Quiñones F. 2003. Bacterial iron homeostasis. FEMS Microbiol Rev 27:215–237. doi:10.1016/s0168-6445(03)00055-x

Aper SJA, Spreeuwel ACC van, Turnhout MC van, Linden AJ van der, Pieters PA, Zon NLL van der, Rambelje SL de la, Bouten CVC, Merkx M. 2014. Colorful Protein-Based Fluorescent Probes for Collagen Imaging. PloS One 9:e114983. doi:10.1371/journal.pone.0114983

Arnison PG, Bibb MJ, Bierbaum G, Bowers AA, Bugni TS, Bulaj G, Camarero JA, Campopiano DJ, Challis GL, Clardy J, Cotter PD, Craik DJ, Dawson M, Dittmann E, Donadio S, Dorrestein PC, Entian K-D, Fischbach MA, Garavelli JS, Göransson U, Gruber CW, Haft DH, Hemscheidt TK, Hertweck C, Hill C, Horswill AR, Jaspars M, Kelly WL, Klinman JP, Kuipers OP, Link AJ, Liu W, Marahiel MA, Mitchell DA, Moll GN, Moore BS, Müller R, Nair SK, Nes IF, Norris GE, Olivera BM, Onaka H, Patchett ML, Piel J, Reaney MJT, Rebuffat S, Ross RP, Sahl H-G, Schmidt EW, Selsted ME, Severinov K, Shen B, Sivonen K, Smith L, Stein T, Süssmuth RD, Tagg JR, Tang G-L, Truman AW, Vederas JC, Walsh CT, Walton JD, Wenzel SC, Willey JM, Donk WA van der. 2012. Ribosomally synthesized and post-translationally modified peptide natural products: overview and recommendations for a universal nomenclature. Nat Prod Rep 30:108–160. doi:10.1039/c2np20085f

Aubert DF, Hamad MA, Valvano MA. 2014. Host-Bacteria Interactions, Methods and Protocols. Methods Mol Biology 1197:311–327. doi:10.1007/978-1-4939-1261-2_18

Bagg A, Neilands JB. 1987. Ferric uptake regulation protein acts as a repressor, employing iron(II) as a cofactor to bind the operator of an iron transport operon in Escherichia coli. Biochemistry-us 26:5471–5477. doi:10.1021/bi00391a039

Bailey TL, Boden M, Buske FA, Frith M, Grant CE, Clementi L, Ren J, Li WW, Noble WS. 2009. MEME Suite: tools for motif discovery and searching. Nucleic Acids Res 37:W202–W208. doi:10.1093/nar/gkp335

Barrett AR, Kang Y, Inamasu KS, Son MS, Vukovich JM, Hoang TT. 2008. Genetic Tools for Allelic Replacement in Burkholderia Species. Appl Environ Microb 74:4498–4508. doi:10.1128/aem.00531-08

Braffman NR, Piscotta FJ, Hauver J, Campbell EA, Link AJ, Darst SA. 2019. Structural mechanism of transcription inhibition by lasso peptides microcin J25 and capistruin. P Natl Acad Sci Usa 116:1273–1278. doi:10.1073/pnas.1817352116

Braun V, Mahren S. 2005. Transmembrane transcriptional control (surface signalling) of the Escherichia coli Fec type. FEMS Microbiol Rev 29:673–684. doi:10.1016/j.femsre.2004.10.001

Braun V, Mahren S, Ogierman M. 2003. Regulation of the FecI-type ECF sigma factor by transmembrane signalling. Curr Opin Microbiol 6:173–180. doi:10.1016/s1369-5274(03)00022-5

Butt AT, Thomas MS. 2017. Iron Acquisition Mechanisms and Their Role in the Virulence of Burkholderia Species. Front Cell Infect Mi 7:460. doi:10.3389/fcimb.2017.00460

Camacho C, Coulouris G, Avagyan V, Ma N, Papadopoulos J, Bealer K, Madden TL. 2009. BLAST+: architecture and applications. BMC Bioinformatics 10:421–421. doi:10.1186/1471-2105-10-421

Cao L, Do T, Link AJ. 2021. Mechanisms of action of ribosomally synthesized and posttranslationally modified peptides (RiPPs). J Ind Microbiol Biot 48:kuab005. doi:10.1093/jimb/kuab005

Cardona ST, Valvano MA. 2005. An expression vector containing a rhamnose-inducible promoter provides tightly regulated gene expression in Burkholderia cenocepacia. Plasmid 54:219–228. doi:10.1016/j.plasmid.2005.03.004

Cheung-Lee WL, Parry ME, Cartagena AJ, Darst SA, Link AJ. 2019. Discovery and structure of the antimicrobial lasso peptide citrocin. J Biol Chem 294:6822–6830. doi:10.1074/jbc.ra118.006494

Cheung-Lee WL, Parry ME, Zong C, Cartagena AJ, Darst SA, Connell ND, Russo R, Link AJ. 2020. Discovery of Ubonodin, an Antimicrobial Lasso Peptide Active against Members of the Burkholderia cepacia Complex. Chembiochem 21:1335–1340. doi:10.1002/cbic.201900707

Chiarini L, Bevivino A, Dalmastri C, Tabacchioni S, Visca P. 2006. Burkholderia cepacia complex species: health hazards and biotechnological potential. Trends Microbiol 14:277–286. doi:10.1016/j.tim.2006.04.006

Coenye T, Vandamme P, Govan JRW, LiPuma JJ. 2001. Taxonomy and Identification of the Burkholderia cepacia Complex. J Clin Microbiol 39:3427–3436. doi:10.1128/jcm.39.10.3427-3436.2001

Darling P, Chan M, Cox AD, Sokol PA. 1998. Siderophore Production by Cystic Fibrosis Isolates of Burkholderia cepacia. Infect Immun 66:874–877. doi:10.1128/iai.66.2.874-877.1998

Delgado MA, Rintoul MR, Fariás RN, Salomón RA. 2001. Escherichia coli RNA Polymerase Is the Target of the Cyclopeptide Antibiotic Microcin J25. J Bacteriol 183:4543–4550. doi:10.1128/jb.183.15.4543-4550.2001

Depoorter E, Canck ED, Peeters C, Wieme AD, Cnockaert M, Zlosnik JEA, LiPuma JJ, Coenye T, Vandamme P. 2020. Burkholderia cepacia Complex Taxon K: Where to Split? Front Microbiol 11:1594. doi:10.3389/fmicb.2020.01594

Dudek C-A, Jahn D. 2021. PRODORIC: state-of-the-art database of prokaryotic gene regulation. Nucleic Acids Res 50:D295–D302. doi:10.1093/nar/gkab1110

Escolar L, Pérez-Martín J, Lorenzo V de. 1999. Opening the Iron Box: Transcriptional Metalloregulation by the Fur Protein. J Bacteriol 181:6223–6229. doi:10.1128/jb.181.20.6223-6229.1999

Figurski DH, Helinski DR. 1979. Replication of an origin-containing derivative of plasmid RK2 dependent on a plasmid function provided in trans. Proc National Acad Sci 76:1648–1652. doi:10.1073/pnas.76.4.1648

Flannagan RS, Linn T, Valvano MA. 2008. A system for the construction of targeted unmarked gene deletions in the genus Burkholderia. Environ Microbiol 10:1652–1660. doi:10.1111/j.1462-2920.2008.01576.x

Ghilarov D, Inaba-Inoue S, Stepien P, Qu F, Michalczyk E, Pakosz Z, Nomura N, Ogasawara S, Walker GC, Rebuffat S, Iwata S, Heddle JG, Beis K. 2021. Molecular mechanism of SbmA, a promiscuous transporter exploited by antimicrobial peptides. Sci Adv 7:eabj5363. doi:10.1126/sciadv.abj5363

Hassett DJ, Howell ML, Ochsner UA, Vasil ML, Johnson Z, Dean GE. 1997. An operon containing fumC and sodA encoding fumarase C and manganese superoxide dismutase is controlled by the ferric uptake regulator in Pseudomonas aeruginosa: fur mutants produce elevated alginate levels. J Bacteriol 179:1452–1459. doi:10.1128/jb.179.5.1452-1459.1997

Hegemann JD, Zimmermann M, Xie X, Marahiel MA. 2015. Lasso Peptides: An Intriguing Class of Bacterial Natural Products. Accounts Chem Res 48:1909–1919. doi:10.1021/acs.accounts.5b00156

Hider RC, Kong X. 2010. Chemistry and biology of siderophores. Nat Prod Rep 27:637–657. doi:10.1039/b906679a

Higgins S, Sanchez-Contreras M, Gualdi S, Pinto-Carbó M, Carlier A, Eberl L. 2017. The Essential Genome of Burkholderia cenocepacia H111. J Bacteriol 199. doi:10.1128/jb.00260-17

Hogan AM, Jeffers KR, Palacios A, Cardona ST. 2021. Improved Dynamic Range of a Rhamnose-Inducible Promoter for Gene Expression in Burkholderia spp. Appl Environ Microb 87:e00647–21. doi:10.1128/aem.00647-21

Hogan AM, Rahman ASMZ, Lightly TJ, Cardona ST. 2019. A Broad-Host-Range CRISPRi Toolkit for Silencing Gene Expression in Burkholderia. Acs Synth Biol 8:2372–2384. doi:10.1021/acssynbio.9b00232

Käll L, Krogh A, Sonnhammer ELL. 2007. Advantages of combined transmembrane topology and signal peptide prediction—the Phobius web server. Nucleic Acids Res 35:W429–W432. doi:10.1093/nar/gkm256

Käll L, Krogh A, Sonnhammer ELL. 2004. A Combined Transmembrane Topology and Signal Peptide Prediction Method. J Mol Biol 338:1027–1036. doi:10.1016/j.jmb.2004.03.016

Knappe TA, Linne U, Zirah S, Rebuffat S, Xie X, Marahiel MA. 2008. Isolation and Structural Characterization of Capistruin, a Lasso Peptide Predicted from the Genome Sequence of Burkholderia thailandensis E264. J Am Chem Soc 130:11446–11454. doi:10.1021/ja802966g

Kodani S, Unno K. 2020. How to harness biosynthetic gene clusters of lasso peptides. J Ind Microbiol Biot 47:703–714. doi:10.1007/s10295-020-02292-6

Koster M, Vossenberg J, Leong J, Weisbeek PJ. 1993. Identification and characterization of the pupB gene encoding an inducible ferric-pseudobactin receptor of Pseudomonas putida WCS358. Mol Microbiol 8:591–601. doi:10.1111/j.1365-2958.1993.tb01603.x

Krewulak KD, Vogel HJ. 2011. TonB or not TonB: is that the question?This paper is one of a selection of papers published in a Special Issue entitled CSBMCB 53rd Annual Meeting Membrane Proteins in Health and Disease, and has undergone the Journals usual peer review process. Biochem Cell Biol 89:87–97. doi:10.1139/o10-141

Kuznedelov K, Semenova E, Knappe TA, Mukhamedyarov D, Srivastava A, Chatterjee S, Ebright RH, Marahiel MA, Severinov K. 2011. The Antibacterial Threaded-lasso Peptide Capistruin Inhibits Bacterial RNA Polymerase. J Mol Biol 412:842–848. doi:10.1016/j.jmb.2011.02.060

Lee ME, DeLoache WC, Cervantes B, Dueber JE. 2015. A Highly Characterized Yeast Toolkit for Modular, Multipart Assembly. Acs Synth Biol 4:975–986. doi:10.1021/sb500366v

Leitão JH, Feliciano JR, Sousa SA, Pita T, Guerreiro SI. 2017. Burkholderia cepacia Complex Infections Among Cystic Fibrosis Patients: Perspectives and ChallengesProgress in Understanding Cystic Fibrosis. doi:10.5772/67712

Li H, Durbin R. 2009. Fast and accurate short read alignment with Burrows–Wheeler transform. Bioinformatics 25:1754–1760. doi:10.1093/bioinformatics/btp324

Li Y, Han Y, Zeng Z, Li W, Feng S, Cao W. 2021. Discovery and Bioactivity of the Novel Lasso Peptide Microcin Y. J Agr Food Chem 69:8758–8767. doi:10.1021/acs.jafc.1c02659

Li Y, Rebuffat S. 2020. The manifold roles of microbial ribosomal peptide–based natural products in physiology and ecology. J Biol Chem 295:34–54. doi:10.1074/jbc.rev119.006545

LiPuma JJ. 2010. The Changing Microbial Epidemiology in Cystic Fibrosis. Clin Microbiol Rev 23:299–323. doi:10.1128/cmr.00068-09

Lorenzo V de, Wee S, Herrero M, Neilands JB. 1987. Operator sequences of the aerobactin operon of plasmid ColV-K30 binding the ferric uptake regulation (fur) repressor. J Bacteriol 169:2624–2630. doi:10.1128/jb.169.6.2624-2630.1987

Love MI, Huber W, Anders S. 2014. Moderated estimation of fold change and dispersion for RNA-seq data with DESeq2. Genome Biol 15:550. doi:10.1186/s13059-014-0550-8

Luscher A, Moynié L, Auguste PS, Bumann D, Mazza L, Pletzer D, Naismith JH, Köhler T. 2018. TonB-Dependent Receptor Repertoire of Pseudomonas aeruginosa for Uptake of Siderophore-Drug Conjugates. Antimicrob Agents Ch 62:e00097–18. doi:10.1128/aac.00097-18

Mahenthiralingam E, Bischof J, Byrne SK, Radomski C, Davies JE, Av-Gay Y, Vandamme P. 2000. DNA-Based Diagnostic Approaches for Identification of Burkholderia cepacia Complex, Burkholderia vietnamiensis, Burkholderia multivorans, Burkholderia stabilis, and Burkholderia cepacia Genomovars I and III. J Clin Microbiol 38:3165–3173. doi:10.1128/jcm.38.9.3165-3173.2000

Mahenthiralingam E, Urban TA, Goldberg JB. 2005. The multifarious, multireplicon Burkholderia cepacia complex. Nat Rev Microbiol 3:144–156. doi:10.1038/nrmicro1085

Maksimov MO, Pan SJ, Link AJ. 2012. Lasso peptides : structure, function, biosynthesis, and engineering. Nat Prod Rep 29:996–1006. doi:10.1039/c2np20070h

Mathavan I, Beis K. 2012. The role of bacterial membrane proteins in the internalization of microcin MccJ25 and MccB17. Biochem Soc T 40:1539–1543. doi:10.1042/bst20120176

Mathavan I, Zirah S, Mehmood S, Choudhury HG, Goulard C, Li Y, Robinson CV, Rebuffat S, Beis K. 2014. Structural basis for hijacking siderophore receptors by antimicrobial lasso peptides. Nat Chem Biol 10:340–342. doi:10.1038/nchembio.1499

Metelev M, Arseniev A, Bushin LB, Kuznedelov K, Artamonova TO, Kondratenko R, Khodorkovskii M, Seyedsayamdost MR, Severinov K. 2017. Acinetodin and Klebsidin, RNA Polymerase Targeting Lasso Peptides Produced by Human Isolates of Acinetobacter gyllenbergii and Klebsiella pneumoniae. Acs Chem Biol 12:814–824. doi:10.1021/acschembio.6b01154

Montalbán-López M, Scott TA, Ramesh S, Rahman IR, Heel AJ van, Viel JH, Bandarian V, Dittmann E, Genilloud O, Goto Y, Burgos MJG, Hill C, Kim S, Koehnke J, Latham JA, Link AJ, Martínez B, Nair SK, Nicolet Y, Rebuffat S, Sahl H-G, Sareen D, Schmidt EW, Schmitt L, Severinov K, Süssmuth RD, Truman AW, Wang H, Weng J-K, Wezel GP van, Zhang Q, Zhong J, Piel J, Mitchell DA, Kuipers OP, Donk WA van der. 2020. New developments in RiPP discovery, enzymology and engineering. Nat Prod Rep 38:130–239. doi:10.1039/d0np00027b

Mott TM, Vijayakumar S, Sbrana E, Endsley JJ, Torres AG. 2015. Characterization of the Burkholderia mallei tonB Mutant and Its Potential as a Backbone Strain for Vaccine Development. Plos Neglect Trop D 9:e0003863. doi:10.1371/journal.pntd.0003863

Noinaj N, Guillier M, Barnard TJ, Buchanan SK. 2010. TonB-Dependent Transporters: Regulation, Structure, and Function. Nat Microbiol 64:43–60. doi:10.1146/annurev.micro.112408.134247

Ochsner UA, Johnson Z, Vasil ML. 2000. Genetics and regulation of two distinct haem-uptake systems, phu and has, in Pseudomonas aeruginosa. Microbiology+ 146:185–198. doi:10.1099/00221287-146-1-185

Ochsner UA, Vasil AI, Vasil ML. 1995. Role of the ferric uptake regulator of Pseudomonas aeruginosa in the regulation of siderophores and exotoxin A expression: purification and activity on iron-regulated promoters. J Bacteriol 177:7194–7201. doi:10.1128/jb.177.24.7194-7201.1995

Pasqua M, Visaggio D, Sciuto AL, Genah S, Banin E, Visca P, Imperi F. 2017. Ferric Uptake Regulator Fur Is Conditionally Essential in Pseudomonas aeruginosa. J Bacteriol 199. doi:10.1128/jb.00472-17

Pinu FR, Villas-Boas SG, Aggio R. 2017. Analysis of Intracellular Metabolites from Microorganisms: Quenching and Extraction Protocols. Metabolites 7:53. doi:10.3390/metabo7040053

Pradenas GA, Myers JN, Torres AG. 2017. Characterization of the Burkholderia cenocepacia TonB Mutant as a Potential Live Attenuated Vaccine. Vaccines 5:33. doi:10.3390/vaccines5040033

Runti G, Ruiz M d CL, Stoilova T, Hussain R, Jennions M, Choudhury HG, Benincasa M, Gennaro R, Beis K, Scocchi M. 2013. Functional Characterization of SbmA, a Bacterial Inner Membrane Transporter Required for Importing the Antimicrobial Peptide Bac7(1-35). J Bacteriol 195:5343–5351. doi:10.1128/jb.00818-13

Salomón RA, Farías RN. 1995. The peptide antibiotic microcin 25 is imported through the TonB pathway and the SbmA protein. J Bacteriol 177:3323–3325. doi:10.1128/jb.177.11.3323-3325.1995

Salomón RA, Farías RN. 1993. The FhuA protein is involved in microcin 25 uptake. J Bacteriol 175:7741–7742. doi:10.1128/jb.175.23.7741-7742.1993

Salomón RA, Farías RN. 1992. Microcin 25, a novel antimicrobial peptide produced by Escherichia coli. J Bacteriol 174:7428–7435. doi:10.1128/jb.174.22.7428-7435.1992

Sass AM, Coenye T. 2020. Low iron-induced small RNA BrrF regulates central metabolism and oxidative stress responses in Burkholderia cenocepacia. PloS One 15:e0236405. doi:10.1371/journal.pone.0236405

Sass AM, Schmerk C, Agnoli K, Norville PJ, Eberl L, Valvano MA, Mahenthiralingam E. 2013. The unexpected discovery of a novel low-oxygen-activated locus for the anoxic persistence of Burkholderia cenocepacia. Isme J 7:1568–1581. doi:10.1038/ismej.2013.36

Sato T, Nonoyama S, Kimura A, Nagata Y, Ohtsubo Y, Tsuda M. 2017. The Small Protein HemP Is a Transcriptional Activator for the Hemin Uptake Operon in Burkholderia multivorans ATCC 17616. Appl Environ Microb 83. doi:10.1128/aem.00479-17

Schäffer S, Hantke K, Braun V. 1985. Nucleotide sequence of the iron regulatory gene fur. Mol Gen Genetics 200:110–113. doi:10.1007/bf00383321

Shastri S, Spiewak HL, Sofoluwe A, Eidsvaag VA, Asghar AH, Pereira T, Bull EH, Butt AT, Thomas MS. 2017. An efficient system for the generation of marked genetic mutants in members of the genus Burkholderia. Plasmid 89:49–56. doi:10.1016/j.plasmid.2016.11.002

Sievers F, Wilm A, Dineen D, Gibson TJ, Karplus K, Li W, Lopez R, McWilliam H, Remmert M, Söding J, Thompson JD, Higgins DG. 2011. Fast, scalable generation of high-quality protein multiple sequence alignments using Clustal Omega. Mol Syst Biol 7:539–539. doi:10.1038/msb.2011.75

Sokol PA, Darling P, Lewenza S, Corbett CR, Kooi CD. 2000. Identification of a Siderophore Receptor Required for Ferric Ornibactin Uptake in Burkholderia cepacia. Infect Immun 68:6554–6560. doi:10.1128/iai.68.12.6554-6560.2000

Stephan H, Freund S, Beck W, Jung G, Meyer J-M, Winkelmann G. 1993. Ornibactins—a new family of siderophores from Pseudomonas. Biometals 6:93–100. doi:10.1007/bf00140109

Taboada B, Estrada K, Ciria R, Merino E. 2018. Operon-mapper: a web server for precise operon identification in bacterial and archaeal genomes. Bioinformatics 34:4118–4120. doi:10.1093/bioinformatics/bty496

Thomas MS. 2007. Iron acquisition mechanisms of the Burkholderia cepacia complex. Biometals 20:431–452. doi:10.1007/s10534-006-9065-4

Tuanyok A, Kim HS, Nierman WC, Yu Y, Dunbar J, Moore RA, Baker P, Tom M, Ling JML, Woods DE. 2005. Genome-wide expression analysis of iron regulation in Burkholderia pseudomallei and Burkholderia mallei using DNA microarrays. FEMS Microbiol Lett 252:327–335. doi:10.1016/j.femsle.2005.09.043

Tyrrell J, Whelan N, Wright C, Sá-Correia I, McClean S, Thomas M, Callaghan M. 2015. Investigation of the multifaceted iron acquisition strategies of Burkholderia cenocepacia. Biometals 28:367–380. doi:10.1007/s10534-015-9840-1

Ussery DW, Kiil K, Lagesen K, Sicheritz-Pontén T, Bohlin J, Wassenaar TM. 2009. The Genus Burkholderia: Analysis of 56 Genomic Sequences. Genome Dyn 6:140–157. doi:10.1159/000235768

Vandamme P, Peeters C. 2014. Time to revisit polyphasic taxonomy. Antonie Van Leeuwenhoek 106:57–65. doi:10.1007/s10482-014-0148-x

Vincent PA, Delgado MA, Farías RN, Salomón RA. 2004. Inhibition of Salmonella enterica serovars by microcin J25. Fems Microbiol Lett 236:103–107. doi:10.1111/j.1574-6968.2004.tb09634.x

Visca P, Leoni L, Wilson MJ, Lamont IL. 2002. Iron transport and regulation, cell signalling and genomics: lessons from Escherichia coli and Pseudomonas. Mol Microbiol 45:1177–1190. doi:10.1046/j.1365-2958.2002.03088.x

Wasserman WW, Sandelin A. 2004. Applied bioinformatics for the identification of regulatory elements. Nat Rev Genet 5:276–287. doi:10.1038/nrg1315

Winsor GL, Khaira B, Rossum TV, Lo R, Whiteside MD, Brinkman FSL. 2008. The Burkholderia Genome Database: facilitating flexible queries and comparative analyses. Bioinformatics 24:2803–2804. doi:10.1093/bioinformatics/btn524

Yu NY, Wagner JR, Laird MR, Melli G, Rey S, Lo R, Dao P, Sahinalp SC, Ester M, Foster LJ, Brinkman FSL. 2010. PSORTb 3.0: improved protein subcellular localization prediction with refined localization subcategories and predictive capabilities for all prokaryotes. Bioinformatics 26:1608–1615. doi:10.1093/bioinformatics/btq249

Yuhara S, Komatsu H, Goto H, Ohtsubo Y, Nagata Y, Tsuda M. 2008. Pleiotropic roles of iron-responsive transcriptional regulator Fur in Burkholderia multivorans. Microbiology 154:1763–1774. doi:10.1099/mic.0.2007/015537-0

Yuzenkova J, Delgado M, Nechaev S, Savalia D, Epshtein V, Artsimovitch I, Mooney RA, Landick R, Farias RN, Salomon R, Severinov K. 2002. Mutations of Bacterial RNA Polymerase Leading to Resistance to Microcin J25*. J Biol Chem 277:50867–50875. doi:10.1074/jbc.m209425200

Zlosnik JEA, Henry DA, Hird TJ, Hickman R, Campbell M, Cabrera A, Chiavegatti GL, Chilvers MA, Sadarangani M. 2020. Epidemiology of Burkholderia Infections in People with Cystic Fibrosis in Canada between 2000 and 2017. Ann Am Thorac Soc 17:1549–1557. doi:10.1513/annalsats.201906-443oc

